# In vivo Assembly of Bacterial Partition Condensates on Circular Supercoiled and Linear DNA

**DOI:** 10.1101/2024.03.26.585537

**Authors:** Hicham Sekkouri Alaoui, Valentin Quèbre, Linda Delimi, Jérôme Rech, Roxanne Debaugny-Diaz, Delphine Labourdette, Manuel Campos, François Cornet, Jean-Charles Walter, Jean-Yves Bouet

## Abstract

In bacteria, faithful DNA segregation of chromosomes and plasmids is mainly mediated by ParABS systems. These systems, consisting of a ParA ATPase, a DNA binding ParB CTPase, and centromere sites *parS*, orchestrate the separation of newly replicated DNA copies and their intracellular positioning. Accurate segregation relies on the assembly of a high-molecular-weight complex, comprising a few hundreds of ParB dimers nucleated from *parS* sites. This complex assembles in a multi-step process and exhibits dynamic liquid-droplet properties. Despite various proposed models, the complete mechanism for partition complex assembly remains elusive. This study investigates the impact of DNA supercoiling on ParB DNA binding profiles *in vivo*, using the ParABS system of the plasmid F. We found that variations in DNA supercoiling does not significantly affect any steps in the assembly of the partition complex. Furthermore, physical modeling, leveraging ChIP-seq data from linear plasmids F, suggests that ParB sliding is restricted to approximately 2-Kbp from *parS*, highlighting the necessity for additional mechanisms beyond ParB sliding over DNA for concentrating ParB into condensates nucleated at *parS*. Lastly, explicit simulations of a polymer coated with bound ParB suggest a dominant role for ParB-ParB interactions in DNA compaction within ParB condensates.

## Introduction

In bacteria, faithful DNA segregation is essential and ensures that cell offsprings receive at least one copy of each replicon, chromosome or plasmids, after their replication. This process involves the separation and transportation of the new copies in opposite directions along the cell’s longitudinal axis (Cornet et al., 2023). The partitioning of chromosomes and most low-copy number plasmids relies on ParABS systems, which consist of ParA, a Walker-type ATPase, and ParB, a site-specific DNA binding protein and CTPase (Bouet and Funnell, 2019; Jalal and Le, 2020). ParB binds to a few centromeric sites, termed *parS*, and further assembles in a large nucleoprotein complex, called the partition complex. ParA action separates the partition complexes, through the stimulation of its ATPase activity by ParB (Ah-Seng et al., 2013), and actively relocates them in opposite directions. The *parS* sites, present in 1-10 copies near the origins of replication, allow the positioning of the Ori domain rapidly after replication by reaching specific locations, which vary depending on the bacterial species and replicons. These positions are either at quarter-cell positions (*e.g.*, P1 and F plasmids (Gordon et al., 1997; Niki and Hiraga, 1997)), the edges of the nucleoid (*e.g., M. xanthus* (Harms et al., 2013)) or the cell poles (*e.g.*, *C. crescentus* (Bowman et al., 2008)), all ensuring an accurate segregation of Ori domains.

Partition complexes are high molecular weight structures. Their assemblies are initiated by a sequential multi-step process (Figure 1): (i) the specific binding of ParB to *parS*, usually a 16-bp DNA motif (Lin and Grossman, 1998; Livny et al., 2007; Pillet et al., 2011), (ii) the binding of CTP to *parS*-bound ParB (Osorio-Valeriano et al., 2019; Soh et al., 2019), followed by (iii) the conversion of ParB into a clamp and its ejection from *parS* due to a steric clash upon ParB remodeling (Soh et al., 2019), (iv) the subsequent diffusion over *parS*-proximal DNA (Soh et al., 2019), and lastly (v) the ParB unloading after clamp opening. Interestingly, it has also been shown that partition complexes are dynamic structures that display some properties of a liquid-droplet, *i.e.,* an assembly that may be mediated by phase separation (Azaldegui et al., 2021). These properties include (i) the rapid exchange of ParB between separate partition complexes in the order of a few minutes (Debaugny et al., 2018; Osorio-Valeriano et al., 2021), (ii) the intracellular mobility of ParB ∼100 times slower inside ParB clusters (concentrated phase) than outside (diluted phase) (Guilhas et al., 2020), (iii) the ability of ParB foci to fuse when ParA is degradated (Guilhas et al., 2020), and (iv) the self-organization of ParB into droplets *in vitro* (Babl et al., 2022). Which mechanisms are at play to assemble these high molecular weight structures, essential for DNA segregation, is not yet fully understood.

**Figure 1:**
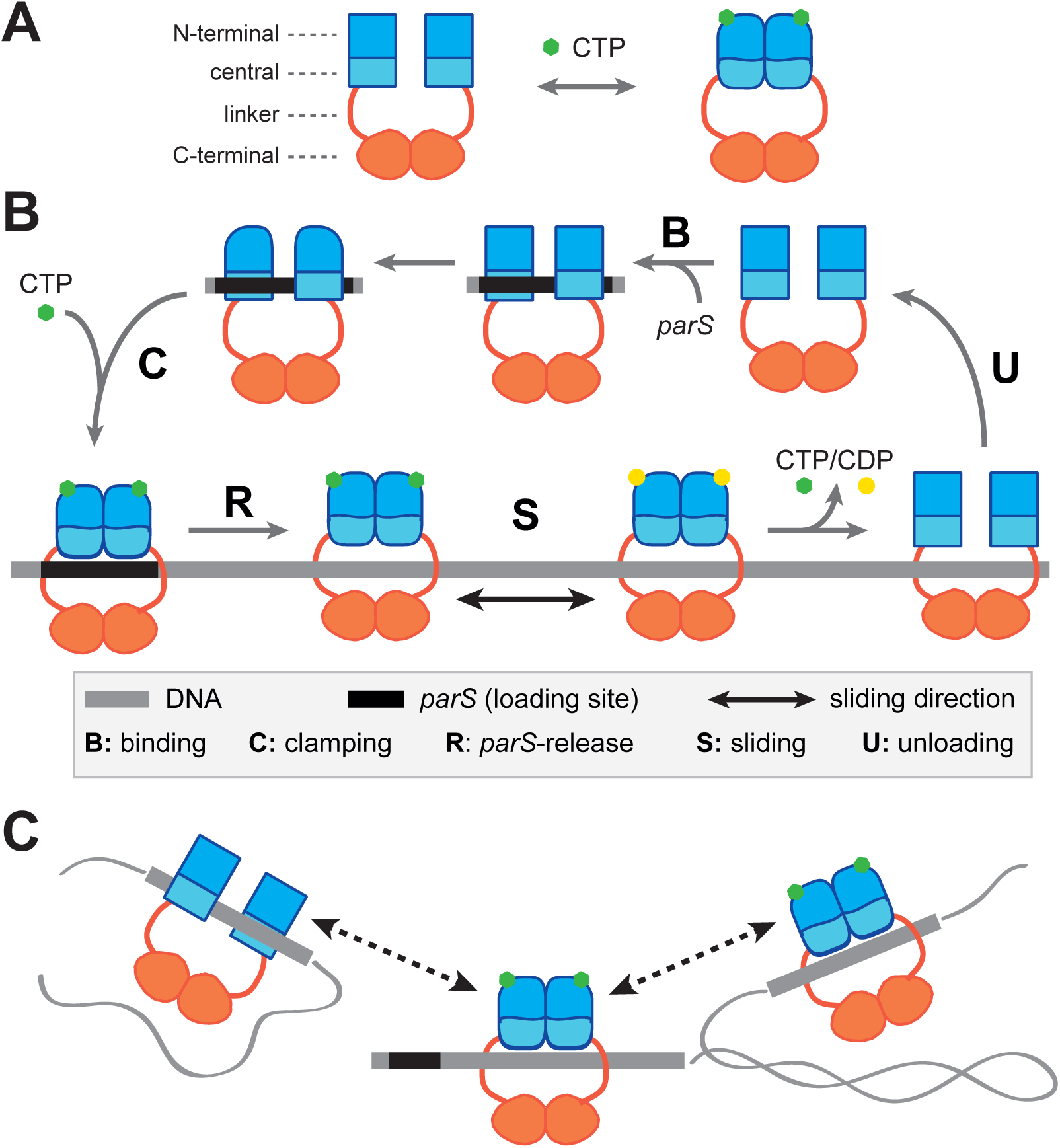
Schematic of ParB dimers and the steps for partition complex assembly. **A-** Open (left) and closed (right) conformations mediated by CTP and *parS* DNA. ParB is a homodimer composed of a C-terminal dimerization domain (orange) link to the central (light blue) and N-terminal (dark blue) domains by a flexible linker (red). The central domain contains the two DNA binding motifs for *parS* binding (Sanchez et al., 2013). The N-terminal part contains the ParA interaction domain, the arginine-like motif, the CTP binding motif and the multimerization domain (Ah-Seng et al., 2009; Soh et al., 2019; Surtees and Funnell, 1999). In the presence of *parS* and CTP, ParB dimer forms a clamp around DNA. **B-** Schematic representation of the initial steps in partition complex assembly. The open conformation of ParB dimer (top right) enables DNA binding to the central part of ParB. Upon specific binding to *parS* centromere (step B), ParB undergoes a conformational change (represented as ParB rounded in the N-terminal part; top left), which promotes CTP binding and subsequently convert ParB as a clamp around *parS* (step C). Clamping promotes ParB release from *parS* by steric clash (Jalal et al., 2021). The ParB clamp “slides” away from *parS* by free diffusion (step S), which allow for a next round of loading at *parS*. After CTP or CDP (upon CTP hydrolysis) unbinding, ParB switches back to an open conformation enabling its unloading from the DNA (step U) hydrolysis. This representation has been updated from (Walter et al., 2020). **C-** Schematic representation of ParB-ParB long-distance interactions arising from ParB diffusing along *parS*-proximal DNA. ParB clamps interact with other ParB dimers, either in the open (left) or close (right) conformations, potentially leading to transient bridging associated with DNA condensation (Balaguer et al., 2021; Tisma et al., 2023).

Several models have been suggested to explain the assembly of the partition complex, including the most recent ones, namely ‘Nucleation & caging’ (N&C) and ‘Clamping & sliding’ (C&S) (Jalal and Le, 2020). This latter was proposed based on the finding that ParB is a CTPase that clamps over *parS* DNA and subsequently diffuses along the DNA until the clamp opens (Figure 1B). The physical modeling of ParB DNA profiles through an exponential decay of ParB binding probability along the *parS*-proximal DNA could depict the ‘C&S’ on naked DNA (Osorio-Valeriano et al., 2021; Walter et al., 2020). However, this model did not align with some biochemical characteristics of ParB, especially the rates of release from *parS* and of unloading from the DNA, thus failing to explain the fast reloading of the partition complex post DNA replication (Walter et al., 2020). Moreover, it does not take into account the presence, *in vivo*, of numerous proteins bound all along the DNA that are expected to impede ParB diffusion (Walter et al., 2020). Conversely, ‘N&C’ describe a long range power-law decay of the probability of ParB binding along the *parS*-proximal DNA which was proposed to account for the attraction of most ParB dimers to a few ParB nucleated from *parS* sites by the combination of low but synergistic interactions, namely ParB-ParB and ParB-non-specific DNA (Debaugny et al., 2018; Sanchez et al., 2015). This stochastic binding model, however, was proposed before the finding that ParB forms CTP-dependent clamps. A hybrid model combining these two frameworks has thus been proposed to describe the ParB DNA binding profile with ‘C&S’ predominantly acting at short distances from *parS*, while ‘N&C’ playing a primary role at longer distance (Walter et al., 2020). However, this model hypothesis has yet to be fully tested.

The modeling of ‘Nucleation & caging’ suggests that DNA compaction directly contributes to the enrichment of ParB around *parS*; the more compact the DNA, the greater the overlaps with ParB condensates (Sanchez et al., 2015). In living systems, DNA compaction predominantly arises from DNA supercoiling (Junier et al., 2023) and/or DNA bridging (van der Valk et al., 2014). In this study, we explored the impact of DNA supercoiling on the ParB DNA binding profiles *in vivo*. Our findings indicate that the assembly of the partition complex of the plasmid F is largely insensitive to significant variations in DNA supercoiling. Moreover, using linear plasmid DNA, we observed an unaltered ParB DNA binding profile compared to supercoiled DNA. Through physical modeling from the ChIP-seq data obtained on a linear plasmid F with DNA ends in the ParB spreading zone, we further demonstrated that the ‘C&S’ alone could not account for the assembly of the partition complex. Additionally, explicit simulations of the ParB enriched region suggests that DNA compaction within ParB condensates primarily arises from ParB-mediated DNA bridging. Altogether, these data provide strong support for a model positing the involvement of two distinct mechanisms for the assembly of partition complexes.

## Results

### ParB DNA binding pattern is invariant to DNA supercoiling variations

To investigate whether one or several steps of the assembly of the partition complex is sensitive to DNA supercoiling, we assayed the DNA binding profile of ParB from the plasmid F *in vivo.* This profile, well described by high-resolution ChIP-sequencing (Debaugny et al., 2018; Sanchez et al., 2015), allows to detect ParB binding to *parS* sites, its release from *parS* upon CTP binding as a clamp over DNA as well as the ParB binding at long distance from *parS* (see Figure 1). The ∼100 kbp plasmid F (F1-10B; Debaugny et al., 2018) was conjugated in two natural hosts, *Escherichia coli* and *Salmonella typhimurium* LT2, that display ∼15 % difference in supercoiling density (σ) (Champion and Higgins, 2007). F1-10B was also introduced in *E. coli topA* and LT2 *gyrB*652 mutants, deficient in DNA topoisomerase I and DNA gyrase activities, and harboring a higher and lower negative supercoiling level, respectively, compare to WT (Champion and Higgins, 2007; Conter et al., 1997; Rovinskiy et al., 2019). F1-10B replicates and segregates faithfully in all these strains.

We first assessed the relative supercoiling density among these four strains by introducing pSAH01, a medium copy-number plasmid of small size (3.4-Kbp). Supercoiling density encompasses both local variations on DNA molecules and variations at the population level, on average. pSAH01, serving as a global sensor, was extracted from exponentially growing cell cultures of *E. coli* at 37°C or LT2 at 30°C. Lowering the temperature reduces the negative supercoiling density (Goldstein and Drlica, 1984), thereby allowing for an expanded range of supercoiling levels to be tested. In addition, the LT2 *gyrB* allele is thermosensitive at 37°C. The topoisomers distributions were measured on agarose gels containing chloroquine, followed by densitometric analyses (Figures 2A and S1A). As expected, pSAH01 was more negatively supercoiled in *E. coli topA* compared to WT with an average shift in the topoisomers distribution of ∼2.2. Comparison between pSAH01 from *E. coli* and LT2 revealed a relative difference of 1.1 topoisomers, consistent with previous results (Champion and Higgins, 2007). Lastly, pSAH01 extracted from LT2 and LT2 *gyrB652* displayed a variation of ∼4.4 topoisomers, as quantified from 2D-gel analyses (Figure 2B).

**Figure 2:**
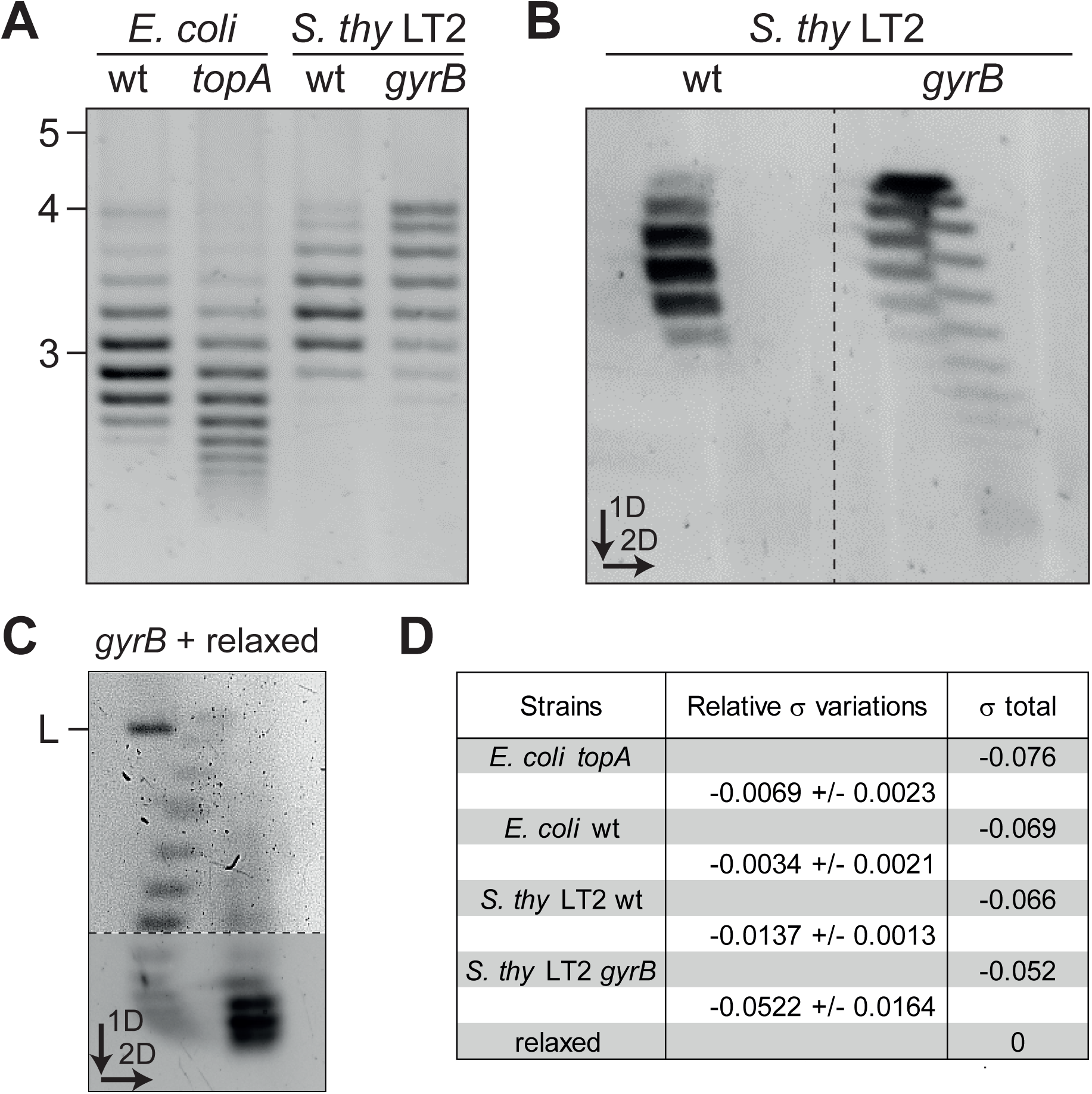
Measurements of DNA supercoiling across various genetic backgrounds. **A-** Relative DNA supercoiling levels in *E. coli*, *S. thy* LT2 and topoisomerase variants *E. coli topA* and *S. thy* LT2 *gyrB* are estimated using the plasmid pSAH01 extracted from growing cultures. *E. coli* and *S. thy* LT2 strains were grown at 37°C and 30°C, respectively. Topoisomers are separated on 1D agarose gel containing chloroquine (5 µg.ml^-1^). **B-** Same as in panel A for *S. thy* LT2 strains resolved in 2D agarose gel containing 5 and 8 µg.ml^-1^ chloroquine in the first and second dimension (indicated by arrows), respectively. **C-** Measurement of total supercoiling density in the *S. thy* LT2 *gyrB* strain. DNA of pSAH01, extracted from *S. thy* LT2 *gyrB*, was separated in two aliquots. One was treated with topoI to relax all supercoils and then mixed back with the untreated aliquot before separation on a 2D agarose gel, as in B with 1.2 and 8 µg.ml^-1^ chloroquine in the first and second dimensions, respectively. The mixing of both samples enables a more accurate calculation of the difference in the number of supercoils (see also Figure S1C). The presence of linear DNA (L) results from plasmid double-strand breaks during the procedures. Note that the top and bottom of the gel are displayed at two different exposures for clarity. Original samples are displayed in Figure S1B. **D-** Summary of relative supercoiling variations and calculated total supercoiling density. The relative supercoiling (σ) variations of pSAH01 between each indicated strain were measured from 1D or 2D agarose gels (described in the main text and Figure S1A). The total DNA supercoiling density is calculated from the relaxed form of pSAH01 (σ = 0) using the relative σ variations between each strain.

As a proxy to estimate the global supercoiling density in each strain, we measured by 2D-gels the variation in topoisomers distribution of pSAH01 between the least negatively supercoiled sample obtained from LT2 *gyrB*652 and its relaxed form (Figures 2C and S1B-C). To note, for an easiest counting of the variation in topoisomers distribution, these two DNA preparations were mixed together (see legends). We found an average shift of 16.9 in the topoisomers distribution between the two samples. Using the formula Δσ = ΔLk / Lk_0_, with Lk_0_ = 323.3 for pSAH01, we calculated an average supercoiling density in LT2 *gyrB652* of −0.052 (Figure 2D). The supercoiling densities in each strain were then estimated using the relative σ variation between them. We found an average of −0.066, −0.069 and −0.076 for LT2 WT, *E. coli* WT and *E. coli topA*, respectively (Figure 2D). Overall, this represents a σ variation of 32% between the two extreme conditions. Considering that, *in vivo*, half of the supercoiling is titrated by proteins bound to DNA (Pettijohn and Pfenninger, 1980), the free supercoiling densities in these strains range from approximately −0.038 to −0.026.

To investigate the impact of DNA supercoiling on the ParB DNA binding profile, we performed ChIP-sequencing assays on the two strains with the greatest difference in σ, *E. coli topA* and LT2 *gyrB*; the former exhibiting ∼50 % more free supercoiled DNA density compare to the latter. The overall ParB_F_ DNA binding profiles were highly reproducible between the two independent duplicates, with ParB_F_ binding detected exclusively on the F plasmid DNA but not on the *E. coli* chromosome, consistent to previous results in another *E. coli* lineage (Figure S2A-B) (Debaugny et al., 2018; Sanchez et al., 2015). Strikingly, the profiles around the *parS*_F_ site from the *topA* and *gyrB*652 strains were nearly identical (Figure 3A). Moreover, the dips and peaks in the signal on the right side of *parS* were essentially observed at the same locations, which is corroborated by a high correlation coefficient (0.988). These data reveal that large intracellular variations in DNA supercoiling levels have no detectable effect on the ParB_F_ DNA binding pattern across the wide range of σ tested. Therefore, this data indicates that none of the steps in the assembly of the partition complex is sensitive to change in DNA supercoiling levels.

**Figure 3:**
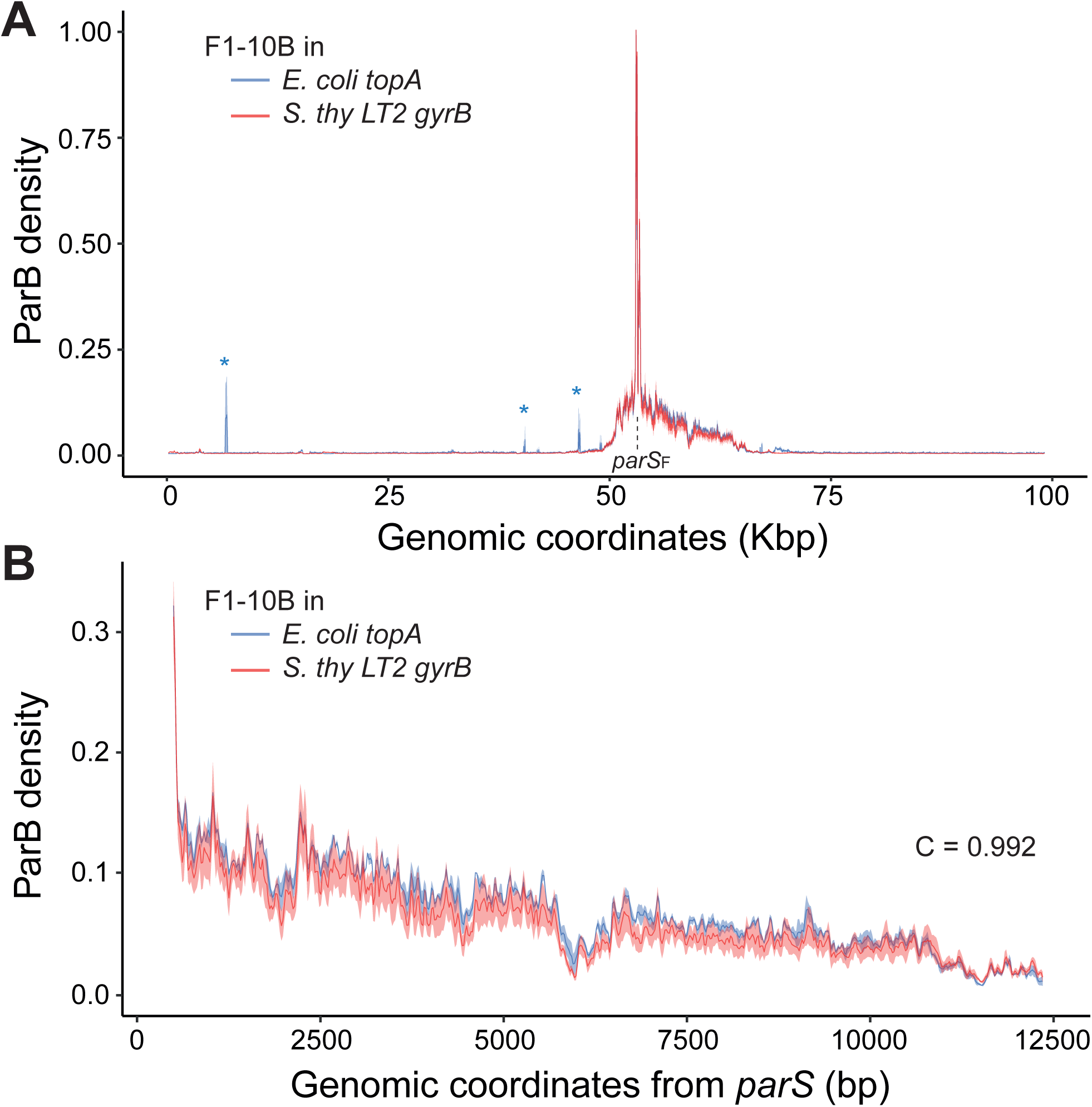
ParB DNA binding profiles are highly similar in the two extreme DNA supercoiling densities. Biological duplicates of ChIP-seq data were performed on *E. coli topA* (blue) and LT2 *gyrB* (red) carrying the F1-10B plasmid. **A-** The ParB reads, normalized to 1 relative to the highest bin, are displayed (ribbon representation) as a function of the genomic coordinates of the plasmid F1-10B, with the line representing the average of each datasets. The *parS*_F_ sites, located between coordinates 53045 and 53447, are indicated by the dashed line. Asterisks represent peaks that are present only in one duplicate of the *topA* dataset (see Figure S2). **B-** Zoom of the data from A on the right side of *parS*, represented as a function of the genomic distance from *parS*. The correlation coefficients (C) are calculated from coordinates 200 to 10200.

### ParB DNA binding patterns on linear DNA molecules

To explore further the sensitivity of the partition complex assembly to DNA supercoiling *in vivo*, we investigated the ParB_F_ DNA binding profile on linear plasmid F DNA. We employed the capability of the coliphage N15 to be maintained as a prophage in a linear plasmid form through the action of the telomerase, TelN, on the *telRL* sites (Ravin, 2003). The cleavage of *telRL* followed by 5’-3’ joining and TelN releasing generates covalently closed hairpin structures at both extremities that are thus protected from exonuclease activities (Deneke et al., 2002). This strategy was previously shown to efficiently linearize the *E. coli* chromosome after insertion of a *telRL* site in the terminus region (Cui et al., 2007). We constructed derivatives of the F1-10B by inserting *telRL* at three positions relative to the *parS* site (3.5-, 13- and 47-Kbp). These F_*telRL* plasmids are circular in the absence of TelN (Figures 4A and S3A-B). When conjugated in a strain carrying the N15 prophage, the F_*telRL* plasmids are efficiently converted to linear DNA molecules as confirmed by a PCR-based assay (Figure S3A-C). DNA supercoils may transiently arise on linear DNA molecules, likely due to local bridging events. However, given the rapid diffusion of DNA supercoiling (Junier et al., 2023), these supercoils should be rapidly eliminated towards the DNA ends, thereby precluding their accumulation.

**Figure 4:**
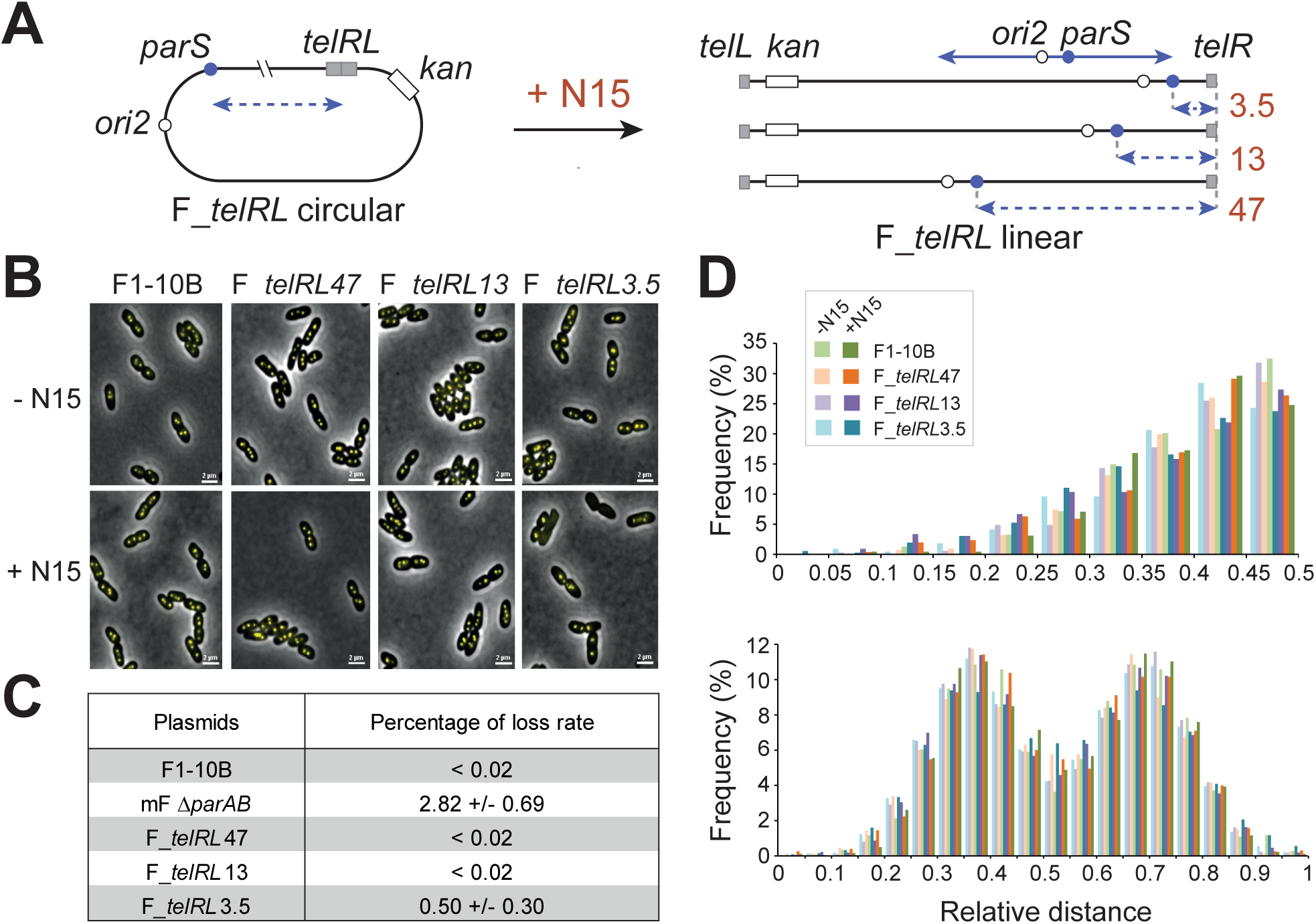
Partition complexes assemble functionally on linear DNA. **A-** Schematic of the plasmid F1-10B with insertions of the *telRL-kan* cassette at 3.5-, 13- and 47-Kbp from *parS*. The open and the blue circles represent the origin of replication (*ori2*) and the centromere site (*parS*), respectively. The dashed blue line represents the variation in distance between *parS* and *telRL* sites. The plasmids F_*telRL* are represented in circular (left) and linear (right) conformations depending on the co-residence of the prophage N15 in the *E. coli* strains (not to scale). **B-** Fluorescence imaging of ParB clusters on circular and linear plasmids *in vivo*. *E. coli* cells carrying (bottom panels) or not (top panels) N15 display foci of ParB_F_-mVenus protein expressed from the endogenous genetic locus on F1-10B and F_*telRL*-mVenus derivatives. Over 99.5% of cells harbor ParB_F_ foci, except for F_*telRL*3.5 in the presence of N15 displaying ∼3% of cells without foci. Scale bars: 2 µm. **C-** Percentage of plasmid loss per generation. F1-10B, F_*telRL* derivatives and mini-F Δ*parAB* were introduced in MC1061/N15 cells, giving rise to linear conformations for the three F_*telRL* derivatives. Standard deviations derived from three independent measurements. **D-** Positioning of the plasmid F1-10B and its derivatives in MC1061, carrying (+) or not (-) the N15 prophage, visualized by fluorescence imaging of ParB-mVenus (see panel B). Statistical analyses of ParB-mVenus foci positioning were performed on cells displaying either one focus (top) or two foci (bottom). The frequency (percentage) of cell with foci located in the indicated intervals is plotted relative to half the cell length (one focus) or the entire cell length (two foci). Lighter and darker colors correspond the absence or presence of N15 prophage as co-resident with F1-10B (green), F_*telRL*47 (red), F_*telRL*13 (violet) and F_*telRL3.5* (blue) plasmids. The number of cells counted with one focus or two foci were 154 to 527 and 1456 to 3228, respectively.

To control that partition complexes are able to assemble on these linear plasmids, we imaged F-*telRL* plasmids, expressing the ParB-mVenus fusion from its endogenous position, by fluorescence microscopy (Figure 4B). We found that, for the three insertion positions, ParB foci are bright with low intracellular background in both the absence and presence of N15. This indicates that partition complexes assemble correctly on the linear F-plasmids. We noticed, however, that in the presence of N15 some cells from the strain carrying F_*telRL*3.5 do not harbor foci, suggesting a defect in plasmid maintenance, either replication or partition. We confirmed this defect by performing a plasmid stability assay (Figure 4C). While the linear plasmids F_*telRL*47 and F_*telRL*13 are as stable as their circular counterparts and the F1-10B (loss rates < 0.02 %), the linear F_*telRL*3.5 is lost at ∼0.5% per cell per generation. This later loss rate, in agreement with the microcopy observation, is however well below random segregation. Indeed, in this growth condition, a circular plasmid F deleted from *parAB* is lost at ∼3%. To discriminate whether this maintenance deficiency arises from a defect in the replication or the partition process, we quantified the intracellular positioning of the F plasmids derivatives, which depend on a functional ParABS system. Indeed, ParB foci, which co-localized with the F-plasmid, are found around mid- or quarter-cell positions in cells with one or two foci, respectively, or are equi-positioned in cells with >2 foci (Diaz et al., 2015). We found that the positions of the three F-*telRL* plasmids, in the circular (-N15) or linear (+N15) forms, are localized around mid-cell (1 focus cells) or quarter positions (two foci cells) similarly to F1-10B (Figure 4D). This indicates that the positioning pattern, and thus partition, is unaffected in the linear forms. The slight loss deficiency observed for F-*telRL*3.5 in the linear form is therefore most likely due to a replication defect. All together, these data indicate that the three linear F plasmids are faithfully segregated, and thus that functional partition complexes are assembled on their *parS* centromere sites independently of the position of the telomerization site.

The assembly of the partition complex on linear F plasmids was further deciphered by investigating their ParB_F_ DNA patterns by ChIP-sequencing, as above. For linear F_*telRL*47 and *F_telRL*13, where the telomerisation sites are located outside of the ParB DNA binding zone, the overall profiles exhibited remarkable similarity, showing high enrichment at and in the close vicinity of *parS* (Figure S4A-B). Notably, when normalized relative to the maximum reads, these ParB DNA binding profiles are also highly similar to those obtained from circular F1-10B plasmids (Figure S5A-B), with the exception of the absence of signal at the *telRL* insertion sites, as expected (insets). In both cases, no change is observed in (i) the initial drop after *parS* sites, (ii) the relative ParB density over the extend of the ParB DNA binding zone and (iii) the pattern of dips and peaks. These data indicate that the linearization of the plasmid F does not alter the specific binding of ParB to *parS*, its ejection upon CTP binding, or its long-range binding over *parS-*proximal DNA, even when a DNA extremity is located only a few Kb from the ParB binding zone. Thus, this confirms that the overall assembly of the partition complex is independent of the global level of DNA supercoiling.

### Partition complex assembly with linearization within the ParB DNA binding zone

We investigated the assembly of the partition complex on the plasmid F linearized at ∼3.5 Kbp of *parS*, *i.e.,* positioning a DNA end in the first part of the ParB DNA binding zone. This is readily observed on the ChIP-seq profiles showing a sharp decline in the number of reads just before the *telRL* site (Figures S4C and S5C). Strikingly, the relative ParB DNA binding profile of F_*telRL*3.5 between *parS* and the telomerization site is very similar to those of circular F1-10B and linear F_*telRL13* and F_*telRL47*. The only distinction lies in the last ∼400-bp, where the signal rapidly drops (Figure 5A). The proximity to the DNA end may account for the gradual reduction in ParB reads to basal level, as observed in the corresponding input within the ∼175-bp proximal to the *telRL*3.5, most likely due to DNA preparation and data processing (Figure S4C, inset). However, unlike the input, the ParB density from the IPs rapidly drops over approximately 225-bp, starting ∼400-bp before the DNA ends and terminating before the last 175-bp. This abrupt drop suggests either a release of clamped ParB due to their escape from the DNA at the free end, a lower probability that the DNA extremity enters in the ParB condensate (boundary effect), or a combination of both.

**Figure 5:**
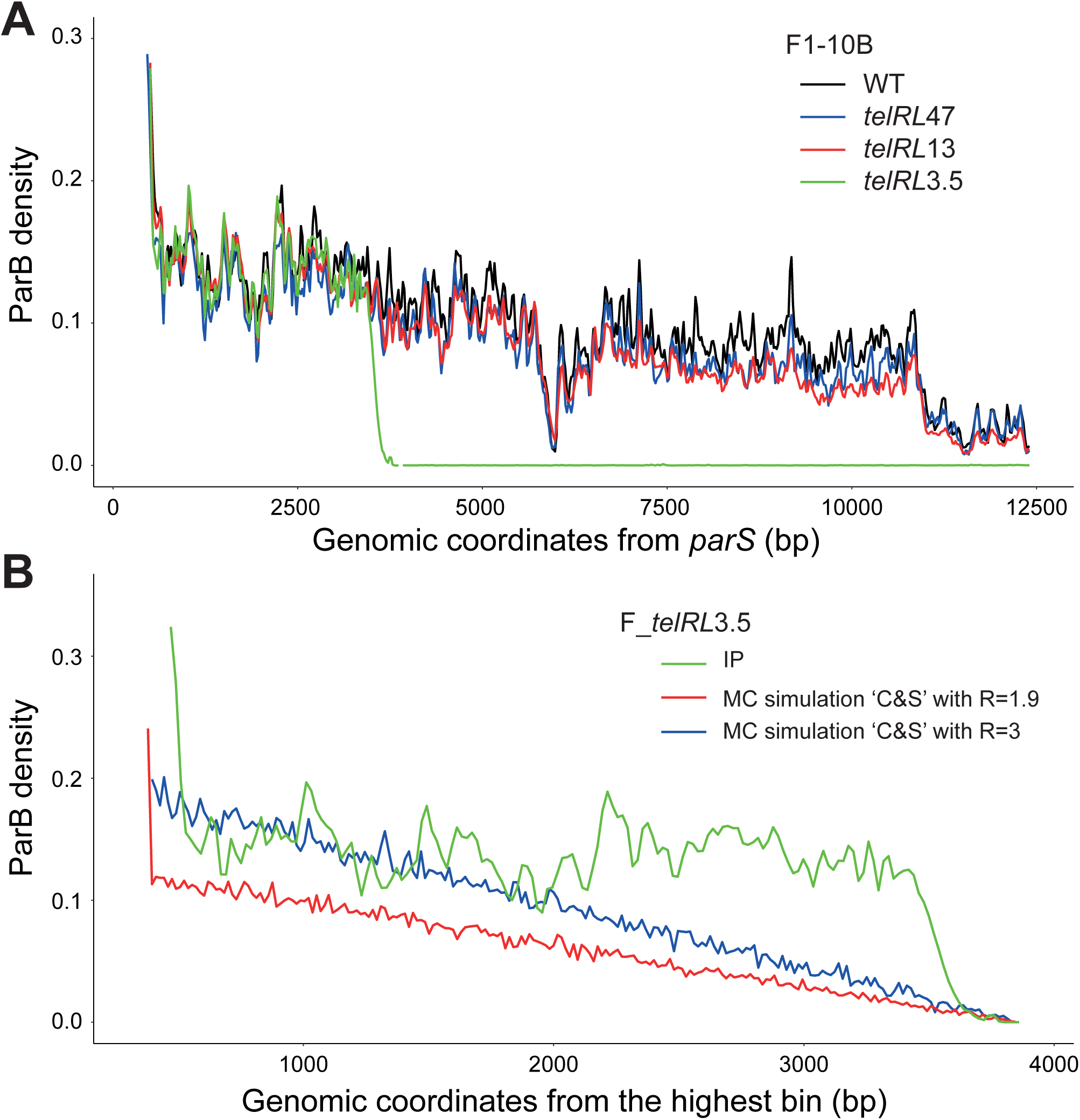
ParB DNA binding profiles is unaffected by linearization of the plasmid F. **A-** Comparison of the ParB DNA binding profiles from ChIP-seq of the circular (wt) and linearized plasmids F1-10B. In the presence of N15, *telRL*47, *telRL*13, *telRL*3.5 are linearized at 47-, 13-, and 3.5-Kbp from *parS*, respectively. The ParB density, relative to the highest bin, is plotted against the genomic coordinates with the origin at the last base of the last *parS* site. Note that for F_*telRL*3.5, the profile ends at the telomerisation site (see Figure S5 for profile on both sides of the telomerisation site). **B-** The ‘Clamping & sliding’ model describes only partly the ParB DNA binding pattern. The ChIP-seq data of the linear F_*telRL*3.5 plasmid (green line), normalized as in panel A, was plotted relative to the coordinate of the highest bin, which is located within the centromere site at about 300-bp from the last *parS* repeat. Monte carlo simulations of the ‘Clamping & sliding’ mechanism have been performed using the same parameters as previously described (see Experimental procedures and Walter et al., 2020), with R, the parameter of ParB release from *parS*, sets as 1.9 or 3 s^-1^ represented by the red and blue lines, respectively. For modeling, the extremity of the linear DNA at which ParB clamps are able to escape was set at 3800-bp from *parS*.

In all cases tested, the relative ParB levels are thus superimposable from *parS* until the drop close the DNA ends (Figure 5A). Moreover, the dips and peaks are at the same locations. This clearly indicates that (i) ParB binding to *parS*, (ii) CTP-dependent ejection of ParB from *parS*, and (iii) subsequent steps, including ParB diffusion over DNA and ParB clustering, remain unchanged between linear and supercoiled DNA.

### ParB clamps diffuse from *parS* over ∼2-Kbp

The ParB DNA binding profile obtained from the linear plasmid F, linearized at 3.5-Kbp from *parS*, provide a valuable data set for assessing whether it can be exclusively described by the ‘Clamping & sliding’ model (Figure 1B). Briefly, the physico-mathematical modeling of this mechanism is based on the equation for the evolution of the unidimensional density described in Experimental procedures (fully described in Walter et al., 2020). In the case of the linear plasmids generated by the TelN-mediated telomerization, which produces uncapped, 5’-3’ hairpin structures at the DNA ends (Deneke et al., 2002), ParB clamps escape from the DNA without the possibility of re-entering from the extremity, as observed *in vitro* (Jalal et al., 2020). Consequently, in the framework of ‘Clamping & sliding’, the DNA ends acts as a sink for ParB clamps. Monte Carlo simulations were then conducted based on this modeling applied to a linear DNA with an extremity (*i.e.*, an absorbing boundary condition (BC) in physical terms) positioned at 3.5-Kbp from *parS* for two sets of parameters.

Firstly, we employed the previously fitted parameters (see experimental procedures), including the release kinetics *R* = 1.9 s^-1^ (Walter et al., 2020). The ‘C&S’ modeling with absorbing BC, in contrast to the periodic BC used for circular plasmids, displays a continuous decrease in ParB density from *parS* to the DNA end where *n*(*x*) =0 (Figure 5B). The boundary effect results in a profile that decreases more rapidly than the periodic BC case (Walter et al., 2020). The ParB DNA binding profile of the linear plasmid F_*telRL*3.5 clearly deviates from the model’s prediction. Indeed, the former shows an identical profile as the circular WT case up to ∼400-bp from the extremity, beyond which the profile drops to zero.

Secondly, given that the overall density of the theoretical profile is diminished compared to previous modeling (Walter et al., 2020) due to the escape of ParB at the telomere, we tested an increased release parameter (*R* = 3 s^-1^) to better align with the experimental data. This adjustment reasonably describes the first half of the ParB density profile but fails to depict accurately the second half. Thus, even with an adjusted release parameter (*R* = 3 s^-1^), the model fits well over a span of ∼2-Kbp, but it also clearly deviates between 2-Kbp and the DNA end. Consequently, this modeling strongly indicates that the ‘Clamping & sliding’mechanism alone cannot entirely account for the ParB binding profile on a DNA with *parS* located at 3.5 Kbp from the DNA end. Another mechanism is therefore required to explain the ParB binding profile after 2-Kbp.

### ParB-ParB interactions dominate supercoiling in DNA compaction within ParB condensates

The invariance of the ChIP-seq ParB profiles following linearization suggests that, within the framework of the ‘N&C’ model (for details, see Figure S6), the average radius of gyration *Rg* of the plasmids remains largely unchanged upon the release of supercoiling. The ‘N&C’ model posits that the primary determinant shaping the ParB profile is the averaged distance of DNA from *parS*, *i.e.*, the radius of gyration (Figure S6A). Supercoiling has been observed to induce approximately 30% DNA compaction (Walter et al., 2021). Based on recent experimental studies (*e.g.*, Balaguer et al., 2021; Tisma et al., 2023) and previous works (*e.g.,* Sanchez et al., 2015; Surtees and Funnell, 1999), we investigated the contribution of *in trans* ParB-ParB interactions to DNA compaction within the ParB enrichment zone.

We performed Monte Carlo simulations (Newman and Barkema, 1999) of a polymer containing interacting particles representing ParB, as depicted in Figure 6A (for details, see Figure S6A). We coarse-grained an open plasmid of 13-Kbp (corresponding to the ParB enriched region) at the experimental resolution scale, *i.e.*, 20-bp (the footprint of ParB binding (Sanchez et al., 2015)), yielding a self-avoiding walk polymer (Vanderzande, 1998) of length N=658 monomers embedded into a face centered cubic (FCC) lattice. Two ParB proteins interact with a strength *J* when they are close in space but distant along DNA (Figure 6A). Initially, *Nb* ∼100 ParB proteins were distributed on the polymer according to the ChIP-seq distribution profile (Figure S7A). Subsequently, the polymer was allowed to move, and after thermalization at the coupling energy *J,* the radius of gyration was sampled every 2.10^4^, corresponding to twice the estimated correlation time. Each sampling can thus be considered as independent. This process was repeated for 1000 realizations of independent ParB distributions.

**Figure 6:**
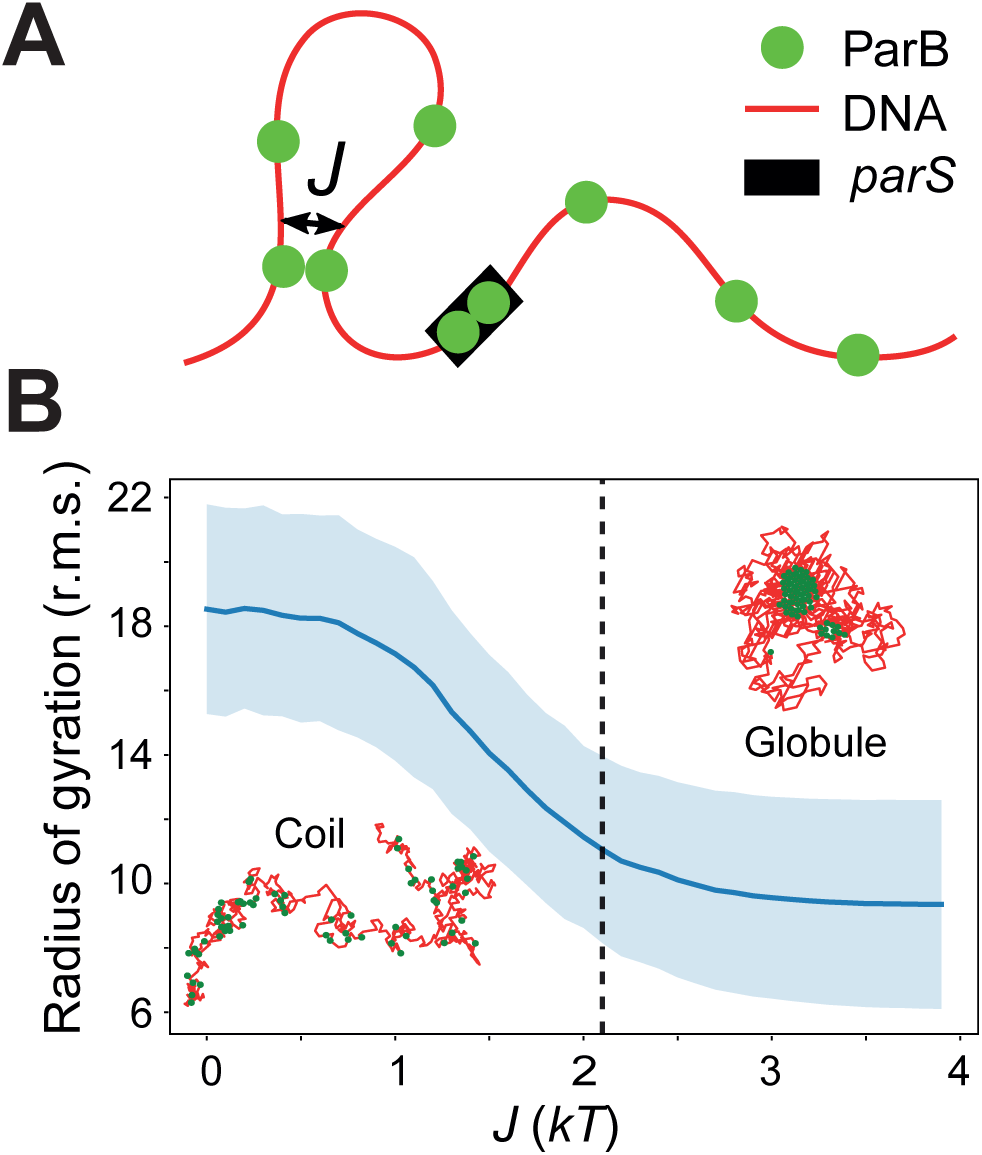
Compaction of DNA induced by ParB-ParB interactions. **A-** Schematic model of the ParB-ParB bridging interactions. The model comprises a coarse-grained DNA polymer (depicted as a red line) of length *L*=658 monomers, each representing 20-bp, corresponding to a total length of ∼13-Kbp, *i.e.,* the region enriched in ParB. ParB proteins (green dots) are distributed according to the ParB DNA binding profile from ChIP-seq assays (see main text). Two close ParB interact with an energy strength *J*. **B**-Dependency of the radius of gyration on ParB-ParB interaction strength *J*. The root mean square (r.m.s.) of the radius of gyration from 1000 independent ParB distributions is plotted versus *J*. The average curve (blue line) displays a transition between a coil and a globule phase at low and high values of *J*, respectively. A drop of ∼50% is observed between the two phases, with a critical transition value of *J_c_* ∼2.1 +/-0.1 *kT* (for details, see Figure S7B).

The simulations were carried out with increasing values of *J* (Figure 6A). The root mean squared radius of gyration plotted against a range of energy *J* shows that the polymer becomes more compact as *J* increases (Figure 6B). Polymers are in coiled conformations when J < 1 and in globular conformations when *J* > 3. We estimated a critical value *J_c_* ∼2.1 +/-0.1 kT (for details, see Figure S7B-E). The level of compaction was computed using the formula (*R_coil_-R_glob_*) / *R_coil_*, where *R_coil_ and R_glob_* represent the radius of gyration in the coil and the globule phases, respectively. We observed a reduction in the radius of gyration by a factor of 48% induced by ParB-ParB interactions. Importantly, the degree of compaction of ∼50% surpasses the ∼30% effect observed with supercoiling (Walter et al., 2021). These modeling data thus corroborate the minimal impact of the variation of DNA supercoiling on DNA compaction within ParB condensates, and support the notion that this compaction primarily arises from ParB-ParB interactions in a supercoiling independent manner.

## Discussion

In this work, we experimentally investigated the role of DNA supercoiling in the assembly of the ParB condensate. Utilizing the conjugative 100-Kbp F-plasmid provided a unique opportunity to test various supercoiling levels in non-isogenic strains and design large linear plasmids that propagates in growing bacterial population. These properties permitted us to analyze ParB_F_ DNA binding profiles over a wide range of supercoiling levels and to demonstrate that DNA supercoiling has minimal, if any, impact on the multi-step assembly of the partition complex. Furthermore, alongside with the ChIP-sequencing data of ParB_F_, physico-mathematical modeling of the DNA binding profiles provides new evidence for a coupled mechanism for explaining ParB binding at long distances from *parS*. It also supports the notion that ParB-ParB bridging interactions, rather than DNA supercoiling, are the primary determinant for DNA compaction in the ParB condensates.

Mutations affecting DNA topology have long been known to induce deficiencies in mini-F plasmid partitioning. In the case of *E. coli gyrB* mutants, such impairment was initially attributed to plasmid relaxation and/or overexpression of ParB_F_, affecting its interaction with *parS* (Ogura et al., 1990). Despite this phenotype, known as IncG incompatibility, has since been elucidated as a consequence of the ParB_F_-induced formation of mini-F multimers in the absence of the ResD/*rfsF* dimer resolution system (Bouet et al., 2006), the question of whether DNA supercoiling influences the assembly of the partition complex remained open. Recent physical modeling, based on ChIP-sequencing data, suggested that DNA compaction needed for the ‘N&C’ based-modeling might be predominantly attributed to DNA supercoiling within the physiological range (Walter et al., 2021). We investigated this possibility by measuring the ParB DNA binding profiles in two natural hosts of the plasmid F mutated for topoisomerase activities, *E. coli topA* and LT2 *gyrB*, exhibiting extreme DNA supercoiling levels (Figure 2). We also investigated F-plasmids propagating as linear DNA molecules and proved to be unaffected in the partitioning process (Figures 4 and S3). We found that ParB DNA binding profiles are highly similar across all conditions, resembling those obtained from the circular plasmid F at wild-type level of total DNA supercoiling. Notably, the release of clamped ParB from *parS*, observed by the rapid decrease of ParB density after *parS* followed by slow decaying over 0.2- to 2-Kbp, is almost superimposable in all conditions tested (Figures 3 and 5). The long distance (2- to 12-Kbp from *parS*) decays of the ParB density are also similar. Thus, our data indicates that none of the steps involved in the auto-assembly of ParB condensates are sensitive to DNA supercoiling level and to the structural form of the DNA molecule, whether circular or linear.

The assembly of partition complex on mini-F plasmids triggers a strong deficit in their negative supercoiling level by 10-12 superhelical turns (Biek and Shi, 1994). This deficit, which is specifically dependent on ParB_F_ and *parS*_F_, has been proposed to arise from interference with the action of the DNA gyrase on the mini-F plasmids (Bouet and Lane, 2009), akin to the ParB silencing that prevents RNA polymerase from accessing promoters in proximity to *parS* (Rodionov et al., 1999). The high local concentration of ParB in the vicinity of *parS*, exceeding 5 mM (Guilhas et al., 2020), would be sufficient to impede the access of DNA gyrase to the mini-F DNA. Indeed, mini-F plasmids typically range between 7- and 12-Kbp in size, a DNA length that could be entirely encompassed within the ParB condensate. In contrast, naturally occurring F plasmids, which range from 70- to 200-Kbp, have a ParB DNA binding zone that represents less than 20 % of the plasmid size, thus allowing DNA gyrase access at long distance from *parS*. The regulation of the homeostasis of DNA supercoiling on such large plasmids should be fully effective given the diffuse nature of supercoils (Junier et al., 2023 and refs therein). One could thus expect that ParB-induced deficit in negative supercoils may not occur, or is strongly reduced, on large, naturally occurring plasmids. This, however, remains to be experimentally determined.

The predominant fraction of intracellular ParB dimers (> 90%) forms densely packed clusters assembled at *parS* sites (Sanchez et al., 2015). These ParB condensates exhibit dynamic characteristics, with a high rate of ParB exchange between condensates (Debaugny et al., 2018; Osorio-Valeriano et al., 2021; Tisma et al., 2023). Upon binding to *parS* and CTP, ParB dimers undergo a significant conformational change, transitioning from *parS* binding to clamping onto flanking DNA (see Figure 1B). Clamped-ParB can diffuse over long distance along naked DNA *in vitro* (Soh et al., 2019). Initially, these properties were considered to be sufficient to explain the ParB DNA binding profiles (Osorio-Valeriano et al., 2021). However, attempts to model these profiles using a ‘Clamping & sliding’ mechanism, based on experimentally determined parameters, failed to explain ChIP-seq data (Walter et al., 2020). Moreover, the presence of numerous roadblocks formed by DNA-bound proteins present on DNA is expected to strongly limit the diffusion of ParB clamps.

In this study, we provided further evidence *in vivo* that the ‘Clamping & sliding’ model alone is not sufficient to explain the ParB DNA binding profile. We took advantage of the linear plasmid F with a telomere site at 3.5-Kbp of *parS*. It displays an unaltered ParB DNA binding profile (Figure 5B), in stark contradiction with the pattern expected from the ‘Clamping & sliding’ model, which predicts a gradual ParB density decay up to the DNA end, acting as a sink for ParB clamps. The ParB density only changes abruptly over ∼225-bp in the last ∼400-bp. Interestingly, however, the ‘Clamping & sliding’ model adequately describes the progressive decay of ParB density up to ∼2-Kbp, suggesting that it can account for the ParB DNA binding profiles only up to a short distance from *parS*. This observation clearly supports the proposal that, in addition to ParB diffusion along the DNA, another mechanism is at play to explain the ParB DNA binding profiles at larger distance from *parS*.

The model ‘Nucleation & caging’ proposed that long-distance ParB-ParB interactions are crucial for condensate formation around *parS* (Sanchez et al., 2015). Recent *in vitro* observations have revealed that ParB can also be loaded independently from *parS* through interaction with a clamped ParB, enabling some ParB to be clamped and diffuse on any DNA present in the spatial proximity to *parS*. This offers the possibility to bypass protein roadblocks and to bridge transiently distant DNA (Tisma et al., 2022). This phenomenon was accounted for in the ‘Nucleation & caging’ model as part of the stochastic ParB-ParB interactions, which also comprise other ParB-ParB interactions, *in cis* and *in trans* of *parS*, not necessarily involving ParB clamps, as observed experimentally (Sanchez et al., 2015; Tisma et al., 2023). In all cases, the combination of all these interactions increases the valency of ParB needed to cluster efficiently most ParB around *parS* and to compact DNA.

The bridging interactions mediated by ParB and CTP have been shown *in vitro* to induce significant compaction specifically on DNA carrying *parS* sequences (Balaguer et al., 2021; Tisma et al., 2023). By employing physical modeling of ParB DNA binding profiles, we assessed the relative contributions of DNA supercoiling and ParB bridging interactions to DNA compaction within ParB condensates. Within the ‘Nucleation & caging’ framework, DNA supercoiling was estimated to yield about 30% DNA compaction *in vivo* (Walter et al., 2021). Employing the same modeling framework, we found that ParB-ParB interactions contribute to DNA compaction by approximately ∼50% (Figure 6B), slightly exceeding the ∼30% attributed to DNA supercoiling. The consistent robustness of the ParB DNA binding profiles in all tested conditions indicates that ParB-ParB interactions acts independently, but not additionally, to DNA supercoiling. Consequently, our results highlight the primary role of the ParB-ParB interactions as the main driver of DNA compaction in ParB condensates. Notably, this mechanism imparts independence to the ParABS system from host factors and enables resilience to various growth conditions. Specifically, for plasmids capable of efficient inter-strain transfer, this independence enables the autonomous assembly of partition condensates independently of the host cells, facilitating adaptation to rapid changes in growth conditions such as environmental stresses affecting DNA supercoiling. Moreover, the presence of highly conserved ParABS systems not only on circular DNA molecules but also on linear plasmids or prophages such as K02, PY54, and N15, as well as on linear chromosomes in bacteria such as *Streptomyces* and *Borrelia*, underscores their adaptability to diverse genetic contexts.

Lastly, it is noteworthy to emphasize that ParB-mediated partition condensates from various plasmid systems serve as thorough and complementary models for understanding bacterial DNA partitioning. Indeed, ParB encoded by plasmids, such as F or RK2, exhibit highly conserved features shared with their chromosomal counterparts. Notably, the binding to dedicated 16-bp *parS* sites and the presence of a ParB DNA binding domain, unusually composed of two distinct centromere binding motifs, called CBM1 and CBM2 (Sanchez et al., 2013). The remarkable conservation of these characteristics may stem from mechanistic constraints, particularly the ejection of ParB clamps arising from the steric hindrance by the CBM2 motif after the CTP-induced remodeling (Soh et al., 2019). Altogether, the mechanisms involved in the assembly of ParB condensates allow for a high level of robustness, which aligns well with the adaptability to nucleate on DNA molecules with varying level of DNA supercoiling, diverse structural shapes and different lengths, whether linear or circular and encompassing plasmids or chromosomes DNA molecules.

## EXPERIMENTAL PROCEDURES

### Bacterial strains, plasmids and oligonucleotides

Strains are derivatives of *E. coli* K12 or *Salmonella typhimurium* LT2 (called LT2 in the text) are listed, together with plasmids, in Table S1. LT2 strains NH2837 and NH2678 are gift from P. Higgins. Constructions of plasmids and strains are detailed in Supplemental experimental procedures. The plasmids F1-10B_*telRL*3.5, F1-10B_*telRL*13 and F1-10B_*telRL*47 were constructed by lambda red recombination from F1-10B through the insertion of a *telRL-kan* cassette at 3.5-, 13- and 47-Kbp from the last repeat of the 16-bp binding motif of *parS*_F_, respectively. Please note that two derivatives of F1-10B_*telRL*3.5, a full-length and a truncated version, were used for ChIP-seq assays (see Supplemental experimental procedures for details). Oligonucleotides used in the study are listed in Table S3.

### Topoisomers analyses

The strains carrying pSAH01 were grown overnight in LB medium, supplemented with ampicillin, under agitation at 37°C or 30°C for *E. coli* or LT2 strains, respectively. The cultures were diluted 200 times in the same medium, and cells were harvested at OD_600_ ∼0.6 by centrifugation of 25 ml. After 2 washes in TE1X, pSAH01 was purified using the MIDI DNA purification kit (Qiagen) and kept à −20°C until use. Topoisomers distributions were analyzed by electrophoresis on 1% agarose gel in 40 mM Tris acetate, 1mM EDTA, as previously described (Bouet and Lane, 2009). Chloroquine concentrations, ranging from 0.5 to 8 µg.ml^-1^ are indicated in the figure legends. Gels were stained with Sybr green before scanning.

Densitometric analyses were performed using Image J software. The relative variation of DNA supercoiling density (Δσ) of pSAH01 between two strains are estimated from the difference in the Gaussian distributions of topoisomers or linking number (ΔLK) divided by the linking number LK_0_ of the plasmid (LK_0_ = plasmid size / 10.5 = 323).

Relaxed CCC DNA was prepared by incubating pSAH01 with Topo I (Invitrogen) in the recommended buffer. Removal of supercoils was verified by agarose gel electrophoresis, and subsequently used in 2D-gel electrophoresis.

### Chromatin immunoprecipitation DNA sequencing (ChIP-seq)

High resolution ChIP-sequencing was carried out using an affinity-purified anti-ParB antibody as previously described (Diaz et al., 2017) with the following modification. Sonication was performed at 10°C using a Covaris® m220 focused ultrasonicator (200 cycles at 75 W for 350 seconds) on aliquots (120 µl), which were then pooled in LoBind tubes for subsequent steps.

The ChIP-seq data (each assays with relevant information are summarized in Table S2) were processed using RStudio software with custom scripts (available upon request). ParB reads were counted at the center of the DNA fragments, taking into account the average size of each DNA library. Background levels on the plasmid F were determined by averaging ParB reads within genomic positions ranging from 1- to 31-Kbp and 74- to 99-Kbp. Following background subtraction, the ParB reads were binned every 20-bp, by applying a smoothing function with averaging over a 40-bp window with a step size of 20-bp. ParB density was obtained by normalizing reads relative to the highest value.

For quantitative comparison between input and IP, the number of reads were normalized relative to the number of mapped reads.

Correlation analyses were performed as previously (Debaugny et al., 2018) using the formula:

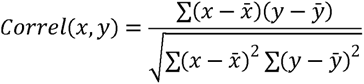

### Epifluorescence microscopy

The strains expressing the fluorescent proteins were grown overnight at 30°C in M9-glucose-CSA medium. The cultures were diluted 250 times in the same medium and incubated at 30°C until OD_600nm_ ∼0.3. 0.7 μl of culture were spotted onto slides coated with a 1% agarose buffered in M9 solution and images were acquired as previously described (Diaz et al., 2015). Nis-Elements AR software (Nikon) was used for image capture and editing. Image analysis was done using ImageJ softwares. The foci counting and positioning on the longitudinal cell axis, were carried out using the macro “Coli inspector” and the plugin “ObjectJ”.

### Plasmid stability assay

The experiments and calculations of the plasmid loss rate per generation were performed essentially as previously described (Sanchez et al., 2013).

### Physical modeling of the ‘Clamping & sliding’ mechanism

The physico-mathematical modeling of ‘Clamping & sliding’ is based on the following equation for the evolution of the unidimensional density *ρ*(*x, t*) of ParB along DNA, fully described in (Walter et al., 2020):

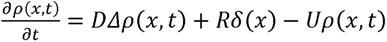

where *x* is the genomic coordinate from *parS*, *t* the time, *D* the diffusion coefficient, *R* the release rate and *U* the unbinding rate. The symbol *Δ* (unidimensional Laplacian) stands for the diffusion (improperly called sliding) and *δ* is the delta function equal to 1 if *x* = 0 and 0 otherwise. It accounts for the release of ParB as a clamp, which occurs exclusively at *parS* (*i.e*., when *x* = 0).

In the stationary state 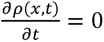, the solution *n*(*x*) = *ρ*(*x, t*)*ε* corresponding to the coarse-grained density of ParB at the scale of the footprint *ε* = 16 *bp* is:

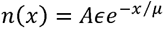

where 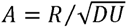 is an overall amplitude and 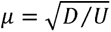 is the characteristic length of the distribution.

The parameters *R*, *D* and *U* have been previously determined with the values *R* = 1.9 s^-1^, *D =* 4.3.10^5^ bp^2^ s^-1^ (∼0.05 µm^2^ s^-1^) and *U* = 4.7.10-3 s^-1^ (Walter et al., 2020).

For the DNA ends close to *parS* (3.5-Kbp), we have used Monte Carlo simulations with the same parameters but with the constraint that the particles disappears from the system when they diffuse across the extremity.

## DATA AVAILABILITY STATEMENT

The raw and processed ChIP-seq data reported in this paper have been deposited in GEO database (NCBI) under the accession number GSE256357. Unpublished custom codes are available upon request.

## ACKNOWLEDGEMENTS

We thank all members of the GeDy (CBI) and the SCPN (L2C) teams for fruitful discussions. We thank P. Higgins for the *S. thy* LT2 strains. We acknowledge the staff at the GeT-Biopuces platform (INSA, Toulouse) for handling libraries and sequencing of the ChIP-seq samples. This work was supported by the CNRS 80Prime MITI grant (Numacoiled) and the CNRS “Modélisation pour le Vivant” MITI Grant (CoilChrom).

## AUTHOR CONTRIBUTIONS

Conceptualization, J.-C.W. and J.-Y.B.;

Data curation, D. L and J.-Y.B.;

Formal analysis, L.D., M.C., J.-Y.B., and J.-C.W.;

Funding Acquisition, J.-C.W. and J.-Y.B.;

Investigation, H.S.A., V.Q., L.D., D.-D.R., J.R., J.-C.W., and J.-Y.B.;

Methodology, H.S.A., V.Q., J.-C.W. and J.-Y.B.;

Project administration, J.-Y.B.;

Resources, F.C., M.C., J.-Y.B.;

Software, M.C., J.-Y.B.;

Supervision, J.-Y.B.;

Validation, H.S.A., J.R., D. L., J.-C.W., and J.-Y.B.;

Visualization, H.S.A., J.-C.W. and J.-Y.B.;

Writing – original draft, H.S.A., J.-C.W. and J.-Y.B.;

Writing – Review & Editing, F.C., H.S.A., J.-C.W., and J.-Y.B.

## ETHICS STATEMENT

This article does not contain any studies with human or animal subjects.

