## Supplementary Material for "In vivo Assembly of Bacterial Partition Condensates on Circular Supercoiled and Linear DNA"

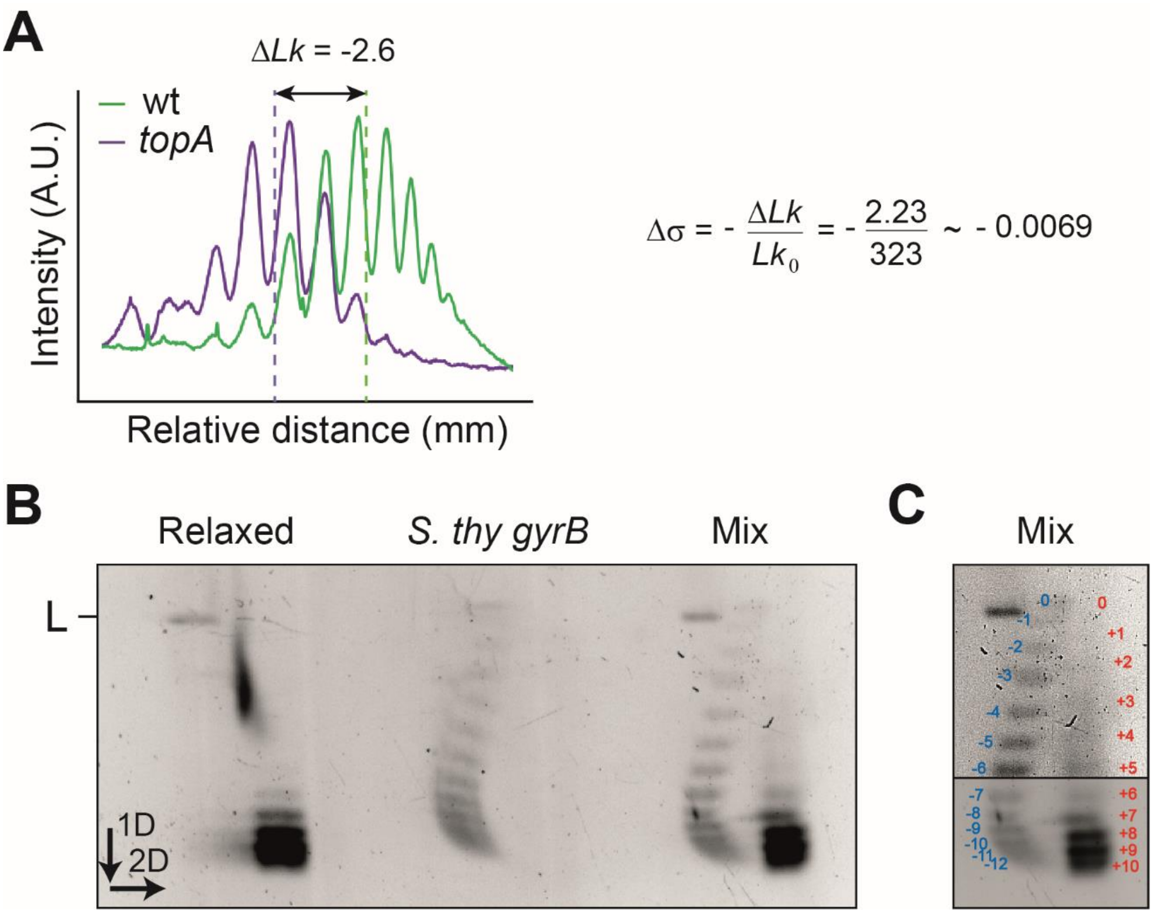


**Figure S1:** Measurements of DNA supercoiling variations between different strains.

**A-** Relative DNA supercoiling variations of pSAH01 extracted from wt and *topA E. coli* strains. (Left) Densitometry profiles, obtained from the same agarose gel (as in Figure 2A), were traced from the wells to align the topoisomers. Note that the intensity is normalized for clarity. The relative variation of linking number ($\Delta Lk$) is determined from the difference in average topoisomers distributions, indicated by the dotted lines (with corresponding colors). (Right) The relative variations in supercoiling density ($\Delta\sigma$) is obtained from the indicated formula with $Lk_0$ corresponding to the linking number in the relax form. $\Delta Lk$ is obtained from three independent biological replicates. **B-** Measurement of the total supercoiling density of pSAH01 in the *S. thy* LT2 *gyrB* strain. pSAH01, extracted from LT2 *gyrB* grown at 30°C, was separated in two aliquots: one (Relaxed) but not the other (LT2 *gyrB*) was treated by topoI to relax supercoils. Samples of each aliquot were also mixed together (Mix). DNA preparations were then subjected to 2D agarose gel electrophoresis in the presence of chloroquine (see legend of Figure 2C). Linear DNA (L), resulting from plasmid double-strand break during the procedures, are indicated. **C-** Same as in B for the mixed DNA preparation, with two different image processing (overexposure in top) to observe all the topoisomers and allow more accurate measurements. Negative and positive supercoils are numbered in blue and red, respectively.

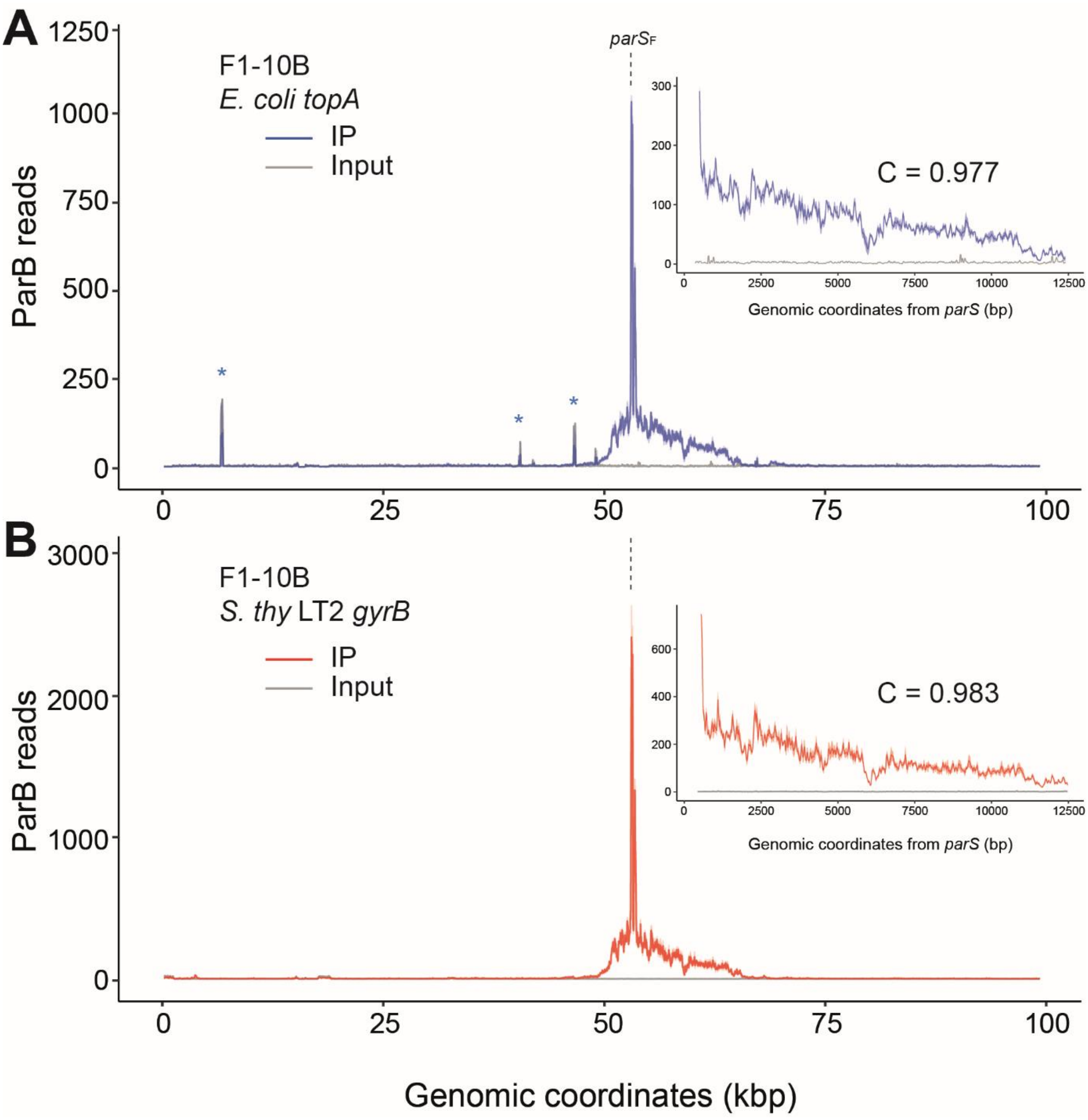


**Figure S2:** ParB DNA binding profiles in *E. coli topA* and LT2 *gyrB* strains.

Biological duplicates of ChIP-seq data were performed on *E. coli topA* (**A**) and LT2 *gyrB* (**B**) carrying the F1-10B plasmid. The normalized ParB reads are displayed (ribbon representation) as a function of the genomic coordinates of the plasmid F1-10B, with the line representing the average of the each datasets. The *parS*$_F$ sites, located between coordinates 53045 and 53447, are indicated by the dashed lines. (Insets) Zoom on the right side of *parS*$_F$. Coordinates are relative to the last bp of the last *parS* site. The correlation coefficients (C) are calculated from coordinates 200 to 10200. Note that experimental variations between duplicates are higher in the *gyrB* than *topA* background. Asterisks in (**A**) represent pics that are present only in one replicate, in both the ChIP and input data (unknown reason).

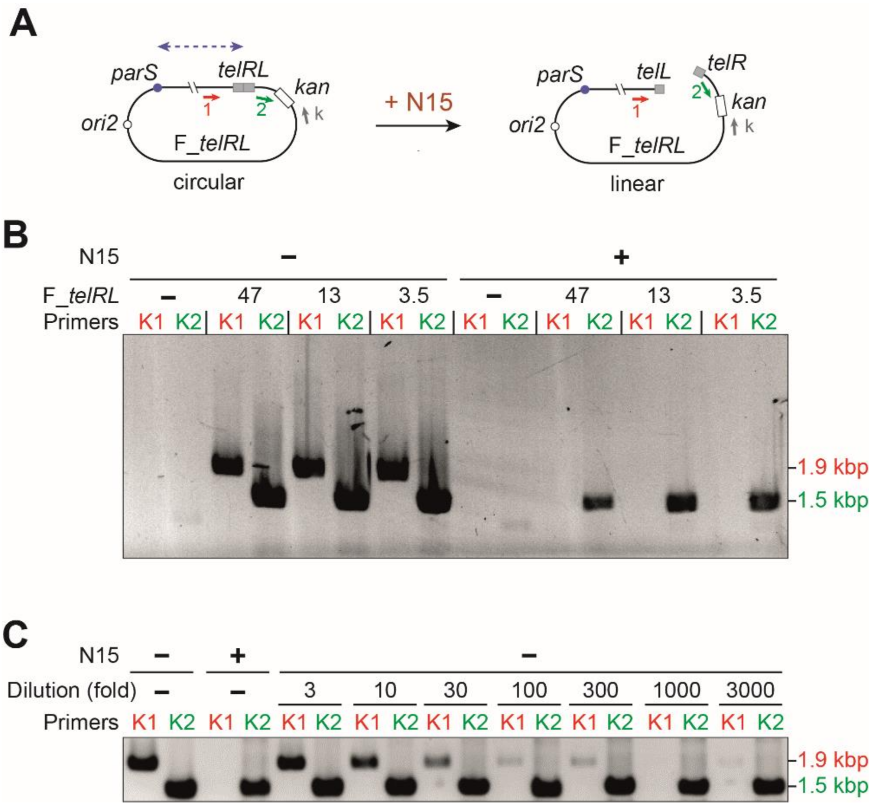


**Figure S3:** Plasmids F1-10B carrying *telRL* insertions are linearized in the presence of N15.

**A-** Schematic of the circular (left) and the linear (right) F_*telRL* plasmids. The *telRL-kan* cassette is inserted at three positions relative to the *parS* site (blue dot), represented by the dashed blue line. In the presence of N15, the telomerase TelN clives at *telRL* and joins the 5' and 3' strands to form covalently closed hairpins ends (for details, see (Ravin, 2003)). Positions of the oligonucleotides, "1", "2" and "k", used to determine the plasmid forms by PCR are indicated, along with their orientations, in red, green and grey, respectively (not to scale). **B-** F_*telRL* plasmids are linear in the presence of N15. PCR, performed on exponentially growing cultures of *E. coli* cells carrying F1-10B derivatives with the indicated insertions of the *telRL* cassette, in the presence (+) or absence (-) of N15, is subjected to agarose gel electrophoresis. In the absence of N15, the two pairs of primers ("k" and "1" (K1) and "k" and "2" (K2); see panel A) produced PCR products of 1.9- and 1.5-kbp indicating that the plasmids are in the circular form. In the presence of N15, only K2 primers give 1.5-kbp PCR products indicating that the plasmids are cleaved by TelN. No PCR products with K1 and K2 primers are observed with the F1-10B wt plasmid, which does not harbor the *telRL-kan* insertion cassette. **C-** All F_*telRL* plasmids are converted in the linear forms in the presence of N15. PCR assays were performed as in panel B from growing cultures of *E. coli* F_*telRL13* cells, carrying (+) or not (-) N15 prophage. The first four lanes done in the absence of dilution and show the linearization in the presence of N15. The sensitivity of the PCR assays to detect circular forms is determined by diluting the culture without N15 (lanes 1-2) with the culture with N15 (lanes 3-4) at the indicated factor. The 1.5-kbp PCR product is thus obtained at nearly the same level for all dilutions. By contrast, the 1.9-kbp product decreases progressively as the dilution factor increases. A faint PCR product is still observed with a dilution of 3000-fold, indicating that the PCR assay would be sensitive enough to detect 1 circular plasmid over 3000 linear one, if present. The absence of the 1.9-kbp PCR product from F_*telRL13* in the presence of N15 (lane 3) indicates that less than 1 circular plasmid over 3000 is present in the population, and thus demonstrates that all plasmids are indeed linear when TelN is produced from N15.

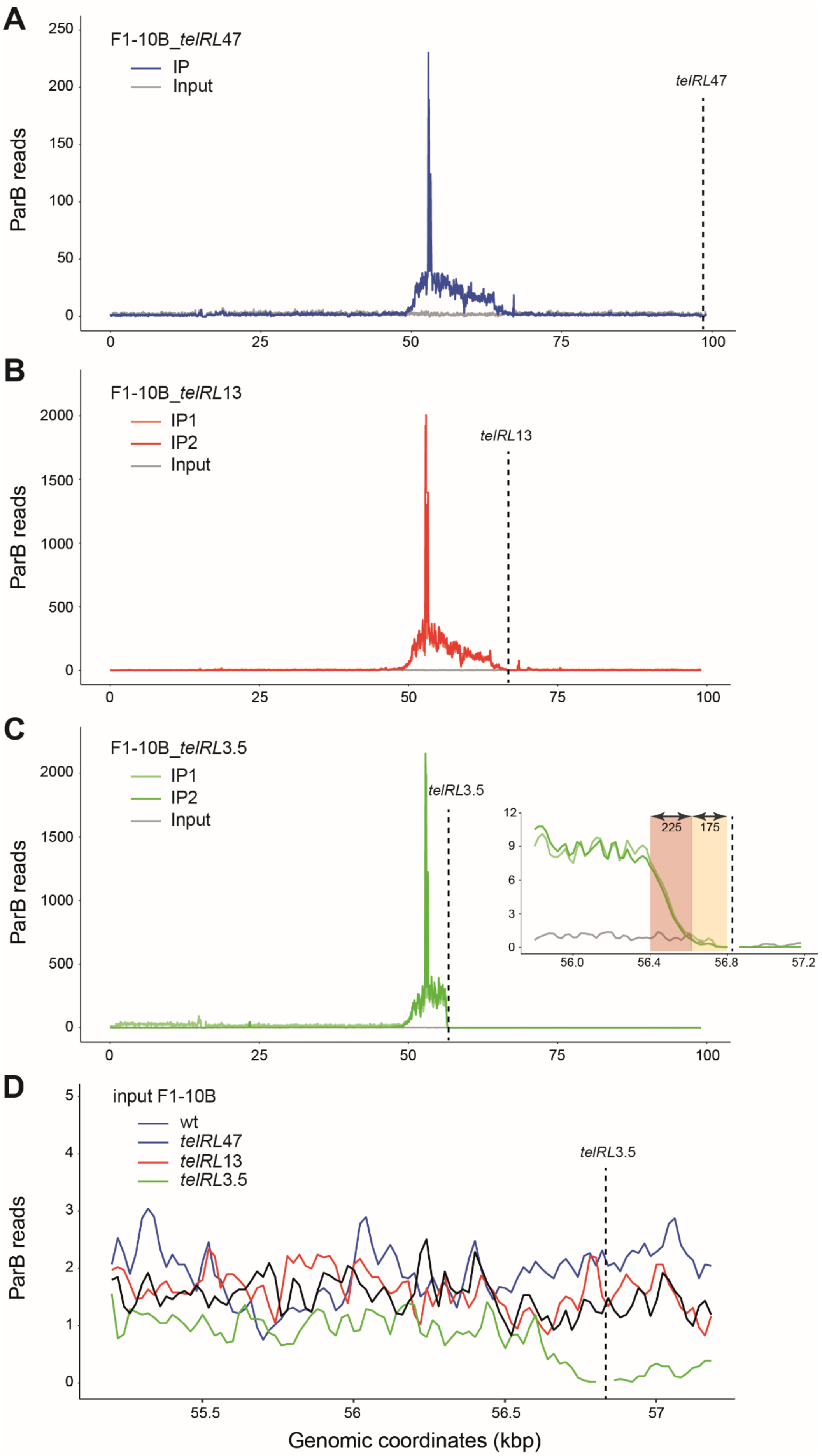

A
F1-10B_telRL47
IP
Input
telRL47
ParB reads
B
F1-10B_telRL13
IP1
IP2
Input
telRL13
ParB reads
C
F1-10B_telRL3.5
IP1
IP2
Input
telRL3.5
225
175
ParB reads
D
input F1-10B
wt
telRL47
telRL13
telRL3.5
telRL3.5
ParB reads
Genomic coordinates (kbp)

**Figure S4:** ParB DNA binding on linear plasmids F1-10B.

ChIP-sequencing assays were performed on the three F_*telRL* plasmids in strains carrying N15. ParB enrichment is only observed on the linear plasmids, not on the *E. coli* chromosome, as for circular F1-10B (Sanchez et al., 2015). The ParB reads (binned) are plotted over the genomic coordinates of F_*telRL*47 **(A)**, F_*telRL*13 **(B)** and F_*telRL*3.5 **(C)**, with the covalently closed hairpin ends indicated by the vertical dotted lines. Inputs correspond to the sequencing of one DNA preparation before IP. IP1 and IP2 represent two biological replicates, normalized relative to the one depicted with the input (grey lines). Note that for F_*telRL*47 only one replicate has been performed. For F_*telRL3.5*, IP2 and IP1 correspond to the full length and truncated (F1-10B_*telRL*3.5t) version of the F1-10B plasmid, respectively. The inset in panel (C) depicts a zoom-in around *telRL*3.5 with the ParB reads of IP1 and IP2 divided by 3.5 and 30 to be displayed alongside the input reads. The colored boxes correspond to the 175-bp (yellow) and 225-bp (ochre), indicated by the doubled arrows, which are proximal to the telRL3.5 site. **(D)** Zoom of the inputs for F1-10B wt (black) and F_*telRL*47 (blue), F_*telRL*13 (red) and F_*telRL*3.5 (green) derivatives between the genomic coordinates 55- and 57.5-kbp.

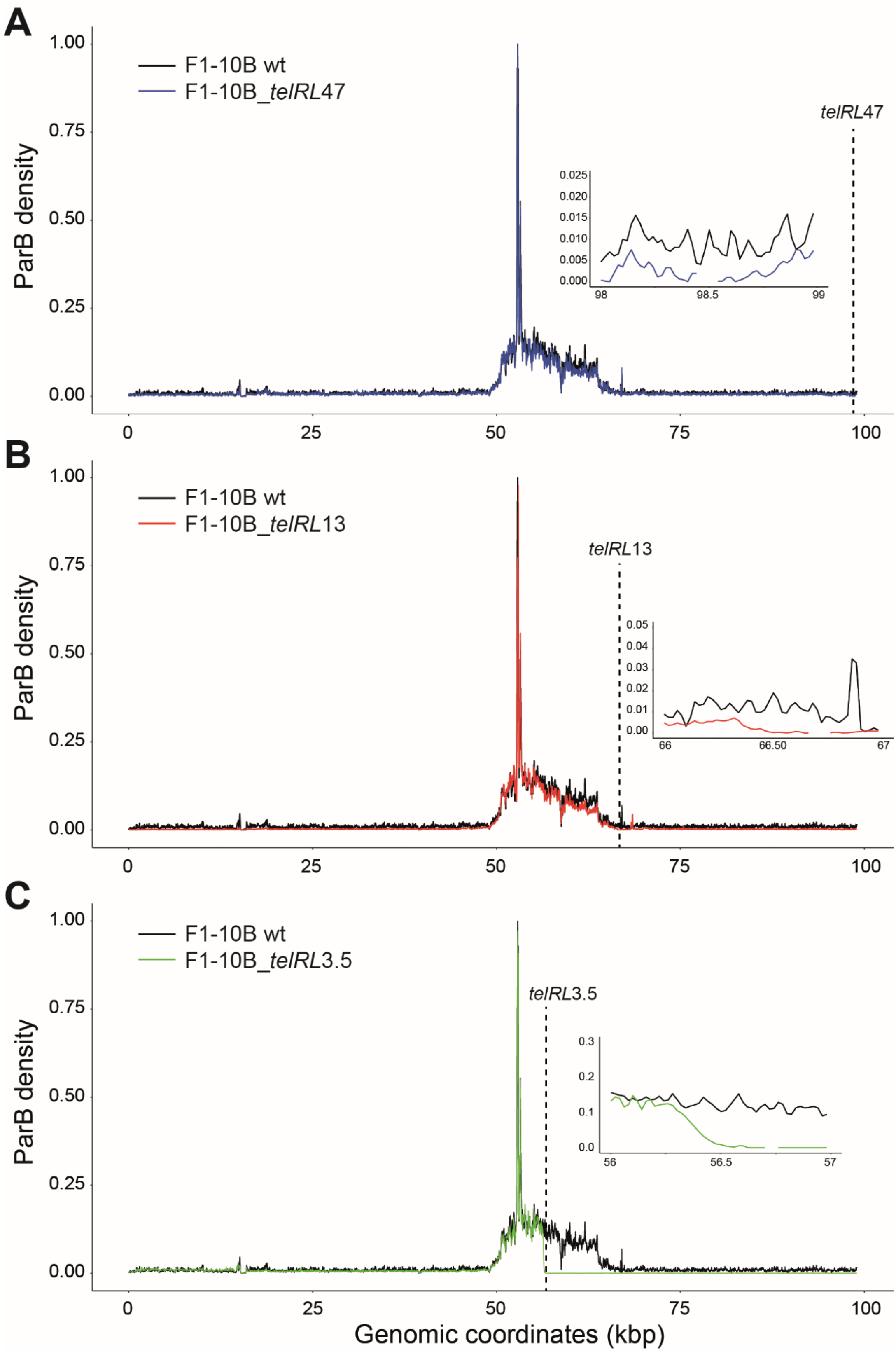


**Figure S5:** ParB DNA binding is highly conserved on linear versus circular plasmids F1-10B. ChIP-seq data (same as in Figure S4) are represented as the average of the duplicates IP1 and IP2 (except for the unique replicate of F_*telRL*47, and between 56.8- to 101.5-kbp for F_*telRL*3.5) density of ParB (1 is the maximum density in a bin), compared to the circular F1-10B (black line). Insets: zoom in the genomic positions of the covalently closed hairpin ends.

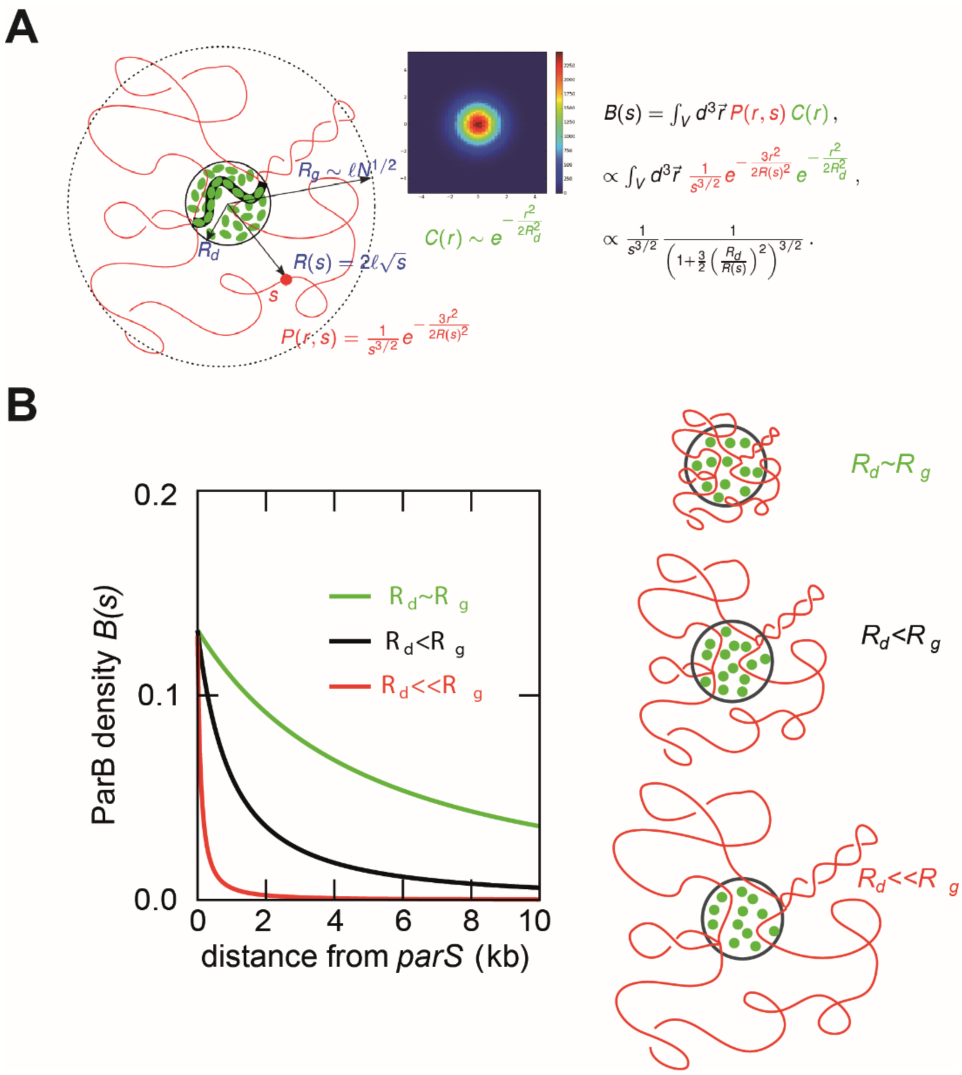


**Figure S6:** The ParB profile predicted by the Stochastic binding model ('Nucleation & caging') is only sensitive to the radius of gyration of DNA. **A-** (*Left*) Schematic of a simple model of Gaussian polymer in interaction with a condensate. The Gaussian polymer is characterized by its length $L = l\ N$, where $l$ is the Kuhn length and $N$ the number of monomers. The average distance $R(s)$ of a locus $s$ from *parS* behaves like $R(s) = 2\ l\ s^{1/2}$ and displays the same scaling behavior as the radius of gyration $Rg \sim l\ N^{1/2}$. The joint probability $P(r, s)$ gives the probability of a locus $s$ to be at the Euclidean distance $r$ from *parS*. The density of the condensate, supposed to be Gaussian too, has a variance $Rd^2$ corresponding, in first approximation, to the averaged radius of the condensate. (*Right*) The theoretical profile $B(s)$ of ParB is readily obtained as the integration of two Gaussian probabilities. The resulting formula displays a dominant term that is a power law of exponent 3/2. The sub-leading correction also depends on the locus $s$ and involves the squared ratio $Rd/R(s)$, *i.e.*, the ratio of the condensate by the averaged distance $r$ to the radius of gyration. **B-** Illustration of the dependency of $B(s)$ versus $Rg$ from three profiles for a plasmid of 10-kbp, with $l$ = 2- (green), 10- (black) and 50-bp (red) corresponding to $Rg$ = 47nm, 104nm and 571nm, respectively. $Rd$ is fixed to 50nm. An overall decrease of the profile is observed as $Rg$ increases, reflecting the decrease of overlap between DNA and the condensate as DNA swells.

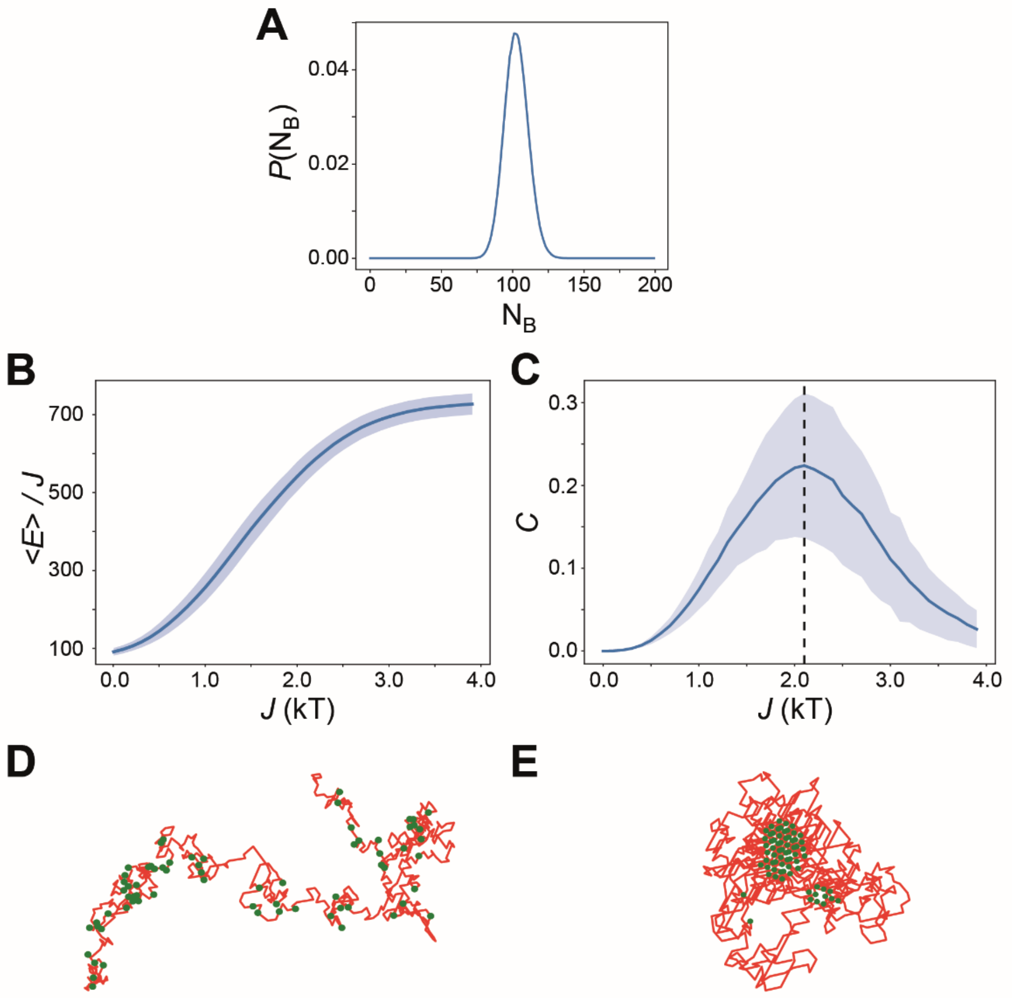


**Figure S7:** Estimation of the critical coupling $Jc$ between a swollen and a compact phase. **A-** Upon an averaging over 1000 independent conformations, the numerical ParB profile is in very good agreement with the conformation of ParB obtained from the ChIP-seq profile of Fig. 3A. We plotted the corresponding probability $P(Nb)$ of having $Nb$ ParB on the enriched region. It displays a Gaussian shape centered around $Nb$=100. **B-** The averaged normalized energy $-\langle E\rangle/J$ is used as the order parameter. The averaging is performed over both thermal fluctuations and initial ParB conformations. The energy of the system is obtained by $E = -\sum_{i,j} \Phi_i U(r_{ij})\Phi_j$, where $\Phi_i$ represents the occupation variable of the monomer $i$ by a ParB protein and takes the values 0 or 1. $U(r_{ij})$ represents the interaction potential between two monomers $i$ and $j$, a function of the Euclidean distance $r_{ij}$ between them. The interaction potential takes the value $J$ when the two monomers are nearest neighbors on the lattice, *i.e.*, $U(r_{ij} = a) = J$ (where $J$ is the energy interaction strength between two proteins), and 0 otherwise. The blue line represents the curve averaged from 1000 independent initial ParB distributions, with the shaded area representing the standard deviation. The polymer is built initially at $J$ = 0 as a self-avoiding polymer. Subsequently, after each increase in the value of $J$, a thermalization with $10^6$ MC steps is performed before the sampling starts. The energy is sampled every $2.10^4$ MC steps, corresponding to twice the calculated correlation time. Each sample can thus be considered as an independent quantity. This curve provides the different phases displayed by the polymer following a change in the interaction energy $J$. Weak ParB-ParB interactions result in a swollen polymer, characterized by a low energy level due to the particle being dispersed. Conversely, strong interactions between particles lead to a compact conformation known as the globule phase, associated with a high energy level due to particles clustering. **C-** The specific heat $C$ is plotted versus $J$ with the same parameters as in (B). The specific heat is defined as $C = (\langle E^2 \rangle - \langle E \rangle^2)/N$ (Newman and Barkema, 1999), and is a precise estimator of the location of the phase transition as it displays a maximum at the critical interaction energy value *Jc* (black dashed line). We estimated *Jc* = 2.1 +/- 0.1$kT$. **D-** Snapshots of a typical polymer conformation ($N$ = 658 monomers; red lines) during a MC simulation at $J$ = 0 (non-interacting particles). Each polymer, containing a number *Nb*~100 of fixed ParB proteins (green dot). The polymer remains swollen as the particles are non-interacting. **E-** Same parameters as (D) but for $J$ = 4, representing strongly interacting particles. The polymer adopts a globular conformation. Due to the sparse distribution of ParB, a compaction of different regions dense in ParB could be observed (2 ParB clusters in this example).

**Supplemental experimental procedures**

**Plasmid constructions**

pSAH01 (3.395-kbp) was constructed from pACYC177 (3.9-kbp) to delete ~0.5-kbp in the *kan* gene by inversed PCR using the PrimeSTAR Max enzyme (Takara Bio) with the primers SAH13 and SAH14 (Table S2). The amplified DNA product, purified using the Gel Band Purification Kit (GE Healthcare), was self-ligated using T4 DNA ligase (New England Biolabs) and transformed in the cloning strain JS238.

The plasmids F1-10B_*telRL*3.5, F1-10B_*telRL*13 and F1-10B_*telRL*47 were constructed by lambda red recombination from F1-10B, or their ParB-mVenus counterparts, through the insertion of a *telRL-kan* cassette at 3.5-, 13- and 47-kbp from the last repeat of the 16-bp binding motif of $parS_F$, respectively. The *telRL-kan* cassette was obtained by PCR amplifications of the *telRL* site from N15 and the *kan* resistance gene from pKD4 with primers carrying extensions of overlapping sequences to allow a subsequent amplification for joining the two PCR products. This cassette was then PCR amplified using the PrimeSTAR Max enzyme (Takara Bio) and the primer pairs RD37.2-RD39, RD40-RD41 and RD42-RD43 for *telRL*3.5, *telRL*13 and *telRL*47, respectively (Table S2). Each of these primers carry an extension of ~30-bp homologous to the targeted insertion sites on the F1-10B plasmids. The gel-purified PCR products *telRL*3.5, *telRL*13, *telRL*47 were dialyzed for 15 min on a 0.025 μm Millipore filter (Merck), transformed by electroporation in strain DY378 carrying F1-10B or F1-10B_BmV and selected for the resistance to kanamycin according to the procedure (Datsenko and Wanner, 2000; Yu et al., 2000). Insertions of the *telRL-kan* cassette were verified by PCR amplification. Note that two versions of F1-10B_*telRL*3.5 have been analyzed by ChIP-seq: a full length (IP2 and input) and a truncated version from coordinates 58.7- to 101.4-Kbp (F1-10B_*telRL*3.5t; ~57-kbp; IP1). They display highly similar profiles between coordinates 0- to 56.8-kbp (see Fig. S4C) and were considered as duplicates.

Plasmids pSAH01 and F1-10B derivatives were introduced in the indicated strains by $CaCl_2$ transformation and conjugation, respectively.

**Table S1:** Bacterial strains and plasmids.

| *Strain* | *Genotype/Relevant properties* | *Source/Reference* |
|---|---|---|
| BLC03 | *S. typhimurium* LT2 wt (NH2837) | Gift from P. Higgins |
| BLC04 | *S. typhimurium* LT2 *gyrB652*$^{ts}$ *zib6794::tn10 ΔTc* (NH2678) | Gift from P. Higgins |
| BLC06 | BLC03 / pSAH01 | This work |
| BLC08 | BLC04 / pSAH01 | This work |
| BLC12 | BLC06 / F1-10B | This work |
| BLC16 | BLC04 / F1-10B | This work |
| BLC17 | BLC08 / F1-10B | This work |
| DLT1215 | W1485 *thi leu thyA deoB supE Δ(ara-leu)7696 zac3051::Tn10 rpsL812* | (Bouet et al., 2007) |
| DLT1370 | W3110 *rpsL topA31::miniTn10* | (Conter et al., 1997) |
| DLT2368 | MC1061 / N15 | Laboratory stock |
| DLT3586 | DLT1215 / F1-10B | This work |
| DLT4028 | DLT1370 / F1-10B | This work |
| DLT4085 | DLT3586 / pSAH01 | This work |
| DLT4092 | MC1061 / F1-10B_*telRL*3.5 | This work |
| DLT4093 | MC1061 / F1-10B_*telRL*13 | This work |
| DLT4094 | MC1061 / F1-10B_*telRL*47 | This work |
| DLT4095 | MC1061 / F1-10B_BmV_*telRL*3.5 | This work |
| DLT4096 | MC1061 / F1-10B_BmV_*telRL*13 | This work |
| DLT4097 | MC1061 / F1-10B_BmV_*telRL*47 | This work |
| DLT4098 | MC1061 / F1-10B | This work |
| DLT4099 | DLT2368 / F1-10B_ *telRL*3.5t | This work |
| DLT4100 | DLT2368 / F1-10B_*telRL*13 | This work |
| DLT4101 | DLT2368 / F1-10B_*telRL*47 | This work |
| DLT4102 | DLT2368 / F1-10B_BmV_*telRL*3.5 | This work |
| DLT4103 | DLT2368 / F1-10B_BmV_*telRL*13 | This work |
| DLT4104 | DLT2368 / F1-10B_BmV_*telRL*47 | This work |
| DLT4105 | DLT2368 / F1-10B | This work |
| DLT4106 | DLT4028 / pSAH01 | This work |
| DLT4121 | DLT2368 / F1-10B_ *telRL*3.5 | This work |
| DLT4164 | DLT2368 / pDAG209 (mF_Δ*parAB*) | This work |
| DLT4166 | MC1061 / pDAG209 (mF_Δ*parAB*) | This work |
| DLT4169 | MC1061 / F1-10B_BmV | This work |
| DLT4170 | DLT2368 / F1-10B_BmV | This work |
| DY378 | *lacI857 Δ(cro-bio)* | (Yu et al., 2000) |
| JS238 | MC1061 *PmalP::lacI*$^{q}$, *srlC::Tn10, recA1* | (Pichoff et al., 1997) |
| MC1061 | MC1061 *araD139 Δ(ara-leu)7679 ΔlacX74 galU galK rpsL thi hsdR2 mcrB* | Laboratory stock |

| *Plasmid* | *Relevant characteristics* | *Source/Reference* |
|---|---|---|
| F1-10B | F1-10 *ccdB cat*$^{+}$ | (Debaugny et al., 2018) |
| F1-10B_*telRL*3.5t | F1-10B *telRL* at 3.5 kb from *parS*, Δ (58- to 101-Kbp) | This work |
| F1-10B_*telRL*3.5 | F1-10B *telRL* at 3.5 kb from *parS* | This work |
| F1-10B_*telRL*13 | F1-10B *telRL* at 13 kb from *parS* | This work |
| F1-10B_*telRL*47 | F1-10B *telRL* at 47 kb from *parS* | This work |
| F1-10B_BmV | F1-10B *parB*$_{F}$-*mVenus* | (Debaugny et al., 2018) |
| F1-10B_BmV_*telRL*3.5 | F1-10B_*telRL*3.5 *parB-mVenus* | This work |
| F1-10B_BmV_*telRL*13 | F1-10B_*telRL*13 *parB-mVenus* | This work |
| F1-10B_BmV_*telRL*47 | F1-10B_*telRL*47 *parB-mVenus* | This work |
| N15 | WT (linear prophage) | (Ravin and Lane, 1999) |
| pDAG209 | mini-F *repFIA*+ *ccdB*- *resD*+ *rsfF*+ *cat*+ Δ*parAB*$_{F}$ | (Bouet et al., 2006) |
| pKD4 | R6K *bla frt-kan-frt* | (Datsenko and Wanner, 2000) |
| pSAH01 | pACYC177 *ampR* Δ*kanR* | This work |

**Table S2**: Synoptic of the information relative to the ChIP-sequencing experiments.

| Strain names | Main properties | Growth condition[a] | ChIP Input/IP Antibody* | Av. Library size** | Reads total | Reads mapped | Repli -cates *** |
|---|---|---|---|---|---|---|---|
| BLC12 | *S. typhimurium* LT2 / F1-10B | 30°C | Input | 149 | 6,554,227 | 6,521,985 | **R1** |
| | | | IP1 antiParB$_F$ | 191 | 9,809,234 | 9,067,162 | **R1** |
| BLC16 | *S. typhimurium* LT2 *gyrB652$^{ts}$* / F1-10B | 30°C | Input | 187 | 5,754,159 | 3,869,340 | **R1** |
| | | | IP1 anti ParB$_F$ | 189 | 10,843,537 | 8,627,520 | **R1** |
| | | | IP2 anti ParB$_F$ | 187 | 1,926,789 | 1,489,543 | **R2** |
| DLT4098 | *E. coli* MC1061 / F1-10B | 37°C | Input | 151 | 5,543,092 | 5,518,379 | **R1** |
| | | | IP anti ParB$_F$ | 162 | 18,110,058 | 18,000,450 | **R1** |
| DLT4099 | *E. coli* MC1061 / N15, F1-10B_telRL3.5 | 37°C | Input | 182 | 4,425,661 | 4,412,379 | **R2** |
| | | | IP anti ParB$_F$ | 173 | 15,852,048 | 1,5493,504 | **R1** |
| | | | | 181 | 12,271,455 | 9,535,898 | **R2** |
| DLT4100 | *E. coli* MC1061 / N15, F1-10B_telRL13 | 37°C | Input | 162 | 6,580,179 | 6,470,513 | **R1** |
| | | | IP anti ParB$_F$ | 164 | 22,526,673 | 21,906,102 | **R1** |
| | | | | 194 | 14,083,917 | 13,693,419 | **R2** |
| DLT4101 | *E. coli* MC1061 / N15, F1-10B_telRL47 | 37°C | Input | 164 | 7,975,041 | 7,835,823 | **R1** |
| | | | IP anti ParB$_F$ | 158 | 23,018,415 | 22,545,774 | **R1** |
| DLT4106 | *E. coli topA* / F1-10B | 37°C | Input | 169 | 11,127,595 | 10,887,278 | **R1** |
| | | | IP1 antiParB$_F$ | 186 | 26,848,805 | 26,075,208 | **R1** |
| | | | IP2 antiParB$_F$ | 187 | 21,217,059 | 20,939,631 | **R2** |

* Affinity-purified antibodies from rabbit-serum.
** Average size of the DNA fragments in the library as measured by sequencing.
*** Replicates indicated in bold correspond to the ones presented in figures in the manuscript.
[a] Cells were grown in LB in exponential phase at the indicated temperature.

**Table S3:** Oligonucleotide sequences.

| *Name* | *Sequence* |
|---|---|
| SAH13 | GGATTCAGTCGTCACTCATGG |
| SAH14 | CATTCTGGTGAAGAAGCTCGACC |
| RD23 | CAGGTTTTCGTCGTCCTCATCGAGCCAGCAGGGGTGATCGCTTTCCTTCACATGGTCCATATGAATATCCTCCTT |
| RD37.2 | TGTTATCGCCGCCTGCGGGCGGCGATACAGGGAGAACGATTAATCATGAAGAATTACGTTGGTATATTTAAAACCTAACTTAATG |
| RD39 | CAGGTTTTCGTCGTCCTCATCGAGCCAGCAGGGGTGATCGCTTTCCTTCACATGGTCCATATGAATATCCTCCTT |
| RD40 | CGGTATCTGACAATGCCTGTAAGATTAATTGTTCTGCACGCTGTTAATTTTTACGTTGGTATATTTAAAACCTAA |
| RD41 | TGCCTGATGTTTATAAATGAGATCTGCCATGCCACTGATTGCTTTGATCTCATGGTCCATATGAATATCCTCCTT |
| RD42 | GTCTTTATTGAAAAACATCAGGCTGAGTTCAGCATCAAAGCAATGTGCCGTTACGTTGGTATATTTAAAACCTAA |
| RD43 | AGAAATATGCTCCGTCACATACCCTTCCGTGTCATACCGGCAGGCACCGGCATGGTCCATATGAATATCCTCCTT |

In Fig. S3, RD37.2, 40 and 42 are termed “1”, RD23 is termed “2” and RD39, 41 and 43 are termed “k”.
